# Prefrontal top-down projections control context-dependent strategy selection

**DOI:** 10.1101/2021.12.14.472559

**Authors:** Olivier Gschwend, Tao Yang, Daniëlle van de Lisdonk, Xian Zhang, Radhashree Sharma, Bo Li

**Affiliations:** Cold Spring Harbor Laboratory, Cold Spring Harbor, NY, USA; Center for Neuroscience, University of Amsterdam, Amsterdam, the Netherlands

## Abstract

The rules governing behavior often vary with behavioral contexts. As a consequence, an action rewarded in one context may be discouraged in another. Animals and humans are capable of switching between behavioral strategies under different contexts and acting adaptively according to the variable rules, a flexibility that is thought to be mediated by the prefrontal cortex (PFC)^1–4^. However, how the PFC orchestrates context-dependent switch of strategies remains unclear. Here we show that pathway-specific projection neurons in the medial PFC (mPFC) differentially contribute to context-instructed strategy selection. In a decision-making task in which mice have been trained to flexibly switch between a previously established rule and a newly learned rule in a context-dependent manner, the activity of mPFC neurons projecting to the dorsomedial striatum encodes the contexts, and further represents decision strategies conforming to the old and new rules. Moreover, the activity of these neuron is required for context-instructed strategy selection. In contrast, the activity of mPFC neurons projecting to the ventral midline thalamus does not discriminate between the contexts, and represents the old rule even if mice have adopted the new one; furthermore, these neurons act to prevent the strategy switch under the new rule. Our results suggest that the mPFC→striatum pathway promotes flexible strategy selection guided by contexts, whereas the mPFC→thalamus pathway favors fixed strategy selection by preserving old rules. Balanced activity between the two pathways may be critical for adaptive behaviors.

## Main text

The ability to rapidly adjust behaviors according to the context is crucial for everyday life. The prefrontal cortex (PFC), a major hub in the brain supporting cognitive flexibility^5–10^, is thought to have a role in context-dependent behavioral responses. For example, the PFC is involved in regulating contextual fear expression^11–14^ and encodes contextual cues that inform rules^2,15,16,6^. In particular, when monkeys engage in a perceptual decision-making task requiring context-guided selection of sensory features (color or motion), PFC neurons represent contextual information in the form of population activity unfolding in a multidimensional space^2,17^, with individual PFC neurons showing mixed selectivity^5,7,18–20^. However, it is unclear whether contextual information in the PFC is essential for context-dependent behavioral responses and, if so, how this information is conveyed to influence such behavioral responses.

The medial PFC (mPFC) projects extensively to the dorsomedial striatum (DMS)^21^, forming a pathway that has been strongly implicated in decision making^22–28^ and behavioral flexibility^28–36^. Another major target of the mPFC is the thalamus. More specifically, the ventral midline thalamic nuclei (VMT) has gained attention in flexible behaviors^37,38^, and the projections from the mPFC to the nucleus reuniens – a part of the VMT – are critical for the formation of contextual fear memories^12^. Previous studies also suggest a role for mPFC-VMT- hippocampal interactions in memory processes^39–41^. Interestingly, recent work indicates that mPFC neurons projecting to the midline thalamus and those projecting to the DMS are both required for flexible behaviors^34^. These findings point to the possibility that the mPFC exerts its functions in flexible behaviors through, at least in part, projections to the DMS and/or VMT.

In this study, we tested the hypothesis that mPFC neurons control context-instructed behavioral flexibility via mPFC→DMS or mPFC→VMT circuit. For this purpose, we first trained mice in an auditory decision-making task based on a two-alternative choice (2AC) task in which animals discriminate “cloud-of-tones” stimuli^42–44^. We further trained the mice to use two contextual cues informing different rules. The first rule required mice to make decisions based on only the sensory evidence in the “clouds” – as they had initially been trained. The second rule required mice to adopt a new decision strategy that relied on both the sensory evidence and reward values. With training, mice learned to make decisions based on the old and new rules in a context-dependent manner, and on a trial-by-trial basis. At different training stages, we imaged the activities of mPFC neurons, including those projecting to the DMS or VMT, and further optogenetically manipulated these neurons while mice performed the task.

We found that mPFC neurons encode, and are essential for, context-guided strategy selection. Notably, through learning, the DMS-projecting (mPFC^DMS^) and VMT-projecting (mPFC^VMT^) neurons acquire opposite functions: whereas mPFC^DMS^ neurons represent contextual changes and are required for switching to the new strategy in a context-dependent manner, mPFC^VMT^ neurons keep a stable representation of the context associated with the initial strategy and impedes the switch. These results uncover distinct roles of mPFC subpopulations in behavioral flexibility and stability.

## mPFC neurons encode and are required for context-dependent decisions

We first trained mice to perform a 2AC task inspired by a previous study^43^. In this task (Fig. 1a), mice initiated a trial by licking the central spout, which was followed by the presentation of a cloud-of-tones stimulus. The “cloud” contained a mixture of tones in a high-frequency (12-17 kHz) or low-frequency (1-6 kHz) range, which were the target tones predicting the delivery of a water reward (3 μl) from the left or right spout, respectively. Thus, the rates of high-frequency tones and low-frequency tones in the cloud determined the strength of sensory evidence for decision making, and hence the difficulty of a trial. The easiest trials had clouds composed of purely the high-frequency or low-frequency target tones (which were 1 and −1 in evidence strength, respectively; Methods), while more difficult trials had clouds consisting of mixed target tones (Fig. 1b). Mice improved performance with training in categorizing different clouds, as suggested by the psychometric function (Fig. 1c, d).

**Fig. 1.**
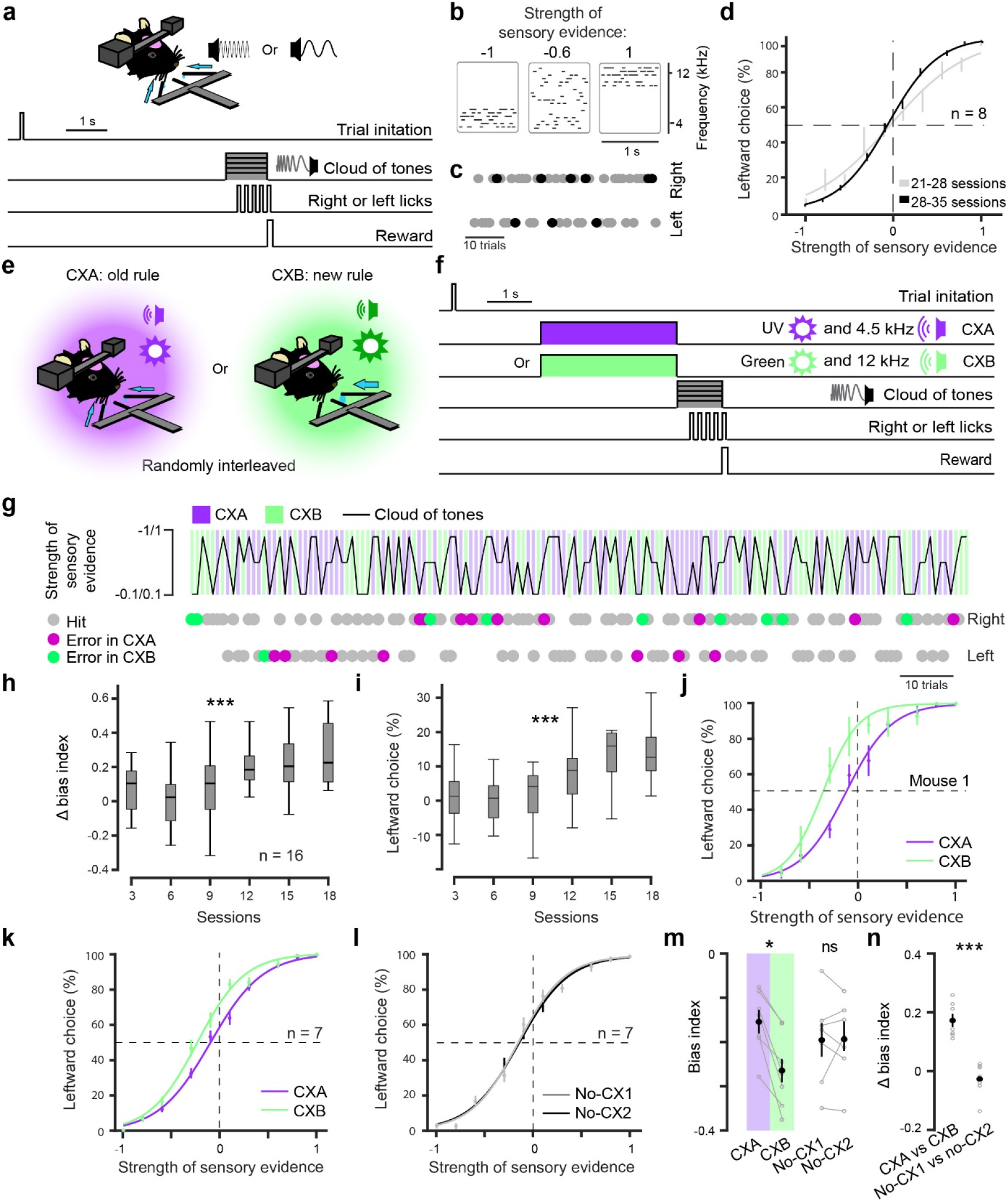
Context-guided two-alternative choice (CX2AC) task. **a,** Schematics of the setup (upper) and structure (lower) of the 2AC task. **b,** Example cloud-of-tones stimuli, in which the strength of sensory evidence for leftward choice is −1 (right), −0.6 (middle), and 1 (left). **c,** Choices in an example session. Gray and black dots represent correct and error trials, respectively. **d,** Psychometric curve of mice (n = 8) averaged across sessions. **e,** Schematics of the contexts and the associated rules in the CX2AC task. **f,** Schematics of the CX2AC task. Note that the CXA and CXB trials were randomly interleaved. **g,** A portion of an example session of the CX2AC task, with upper panel showing the trial-by-trial arrangement of the contexts and the cloud-of-tones stimuli (which are randomly interleaved across trials), and lower panel showing animal choices (left or right) in the corresponding trials. **h,** Changes in bias index across sessions (see Methods; n = 16 mice, F(1,15) = 7.21, ***P = 1.06e-5, one-way ANOVA). **i,** Leftward choice percentage when sensory evidence strength was 0 (F(1,15) = 7.51, ***P = 6.2e-6, one-way ANOVA). **(j, k)** Psychometric curves of an example mouse (j) and 7 mice (k). **l,** Psychometric curves of the same mice as those in (k) in the absence of contextual cues, plotted for two different sets of sessions (no-CX1 & no-CX2). **m,** Bias index calculated from the psychometric curves in (k) and (l) (*P = 0.0378, ns (nonsignificant), P = 0.98, paired Student t-test). **n,** Changes in bias index calculated from the curve in (k) and (l) (***P = 0.00058, paired Student t-test). Psychometric curves are averaged across six sessions. Data are presented as mean ± s.e.m.

Next, we further trained the mice to use contextual information to guide the switch between decision strategies. We added two distinct “contexts” comprising multisensory cues to the 2AC task (Fig. 1e, f; Methods). In each trial, one of the contexts was presented immediately before the onset of the cloud to indicate that one of two rules would be applied. Context A (CXA) informed that the original rule (the “old rule”) would be in effect, so that the mice should keep making choices based on the sensory evidence in the subsequent cloud. Context B (CXB) informed that a large reward (10 μl) would be delivered if mice made a correct leftward choice according to the old rule; however, no rightward choice would lead to any reward. Thus, under this “new rule”, mice still need to use the sensory evidence in the clouds in order to make correct leftward choices, but should ignore the clouds indicating a rightward choice under the old rule. We randomly interleaved CXA trials and CXB trials, but kept CXA trials as the majority (Fig. 1e-g; Methods).

Through training with the contexts, mice gradually adapted their choice strategies to the different rules, as indicated by the increasing bias towards the left spout in CXB trials (Fig. 1h, i). This led to a shift of the psychometric curve towards the left in CXB trials compared with CXA trials, particularly in trials where sensory evidence in the cloud was weak (Fig. 1j, k; Extended Data Fig. 1a). This leftward bias disappeared when the contextual cues were removed (Fig. 1l-n). It is somewhat surprising that the mice still made rightward choices under CXB, especially in trials where the sensory evidence for a rightward choice was strong under the old rule. This is likely because the mice were over trained with the old rule, and the old rule remained in effect in the majority of trials (Methods). Such situations may cause the old rule to interfere with the new rule, an effect that has been previously reported^45^. Taken together, the CX2AC task revealed that mice adaptively select their decision strategies established on the basis of specific action-outcome contingencies (or rules). Importantly, this selection can be flexibly instructed by contextual cues on a trial-by-trial basis.

Previous studies suggest that the dorsolateral PFC in monkeys is involved in context-guided strategy switching^2,46^, and the mPFC in mice controls cognitive flexibility^33,47,48^ and contextual fear learning^11^. To understand how mPFC neurons are recruited and contribute to context-guided strategy selection, we imaged the activity of these neurons in mice performing the CX2AC task. To this end, we injected the mPFC in mice with an adeno-associated virus (AAV) expressing the calcium indicator GCaMP6f^49^, and implanted a cannula into the same location (Fig. 2a, b; Extended Data Fig. 1b-d). We trained the mice in the CX2AC task until they reached a stable performance (Fig. 2c; Extended Data Fig. 1e). At different training stages, we used a wide-field microscope to image the GCaMP6 signals in mPFC neurons at cellular resolution through the implanted cannula^50,51^ (Fig. 2a-c).

**Fig. 2.**
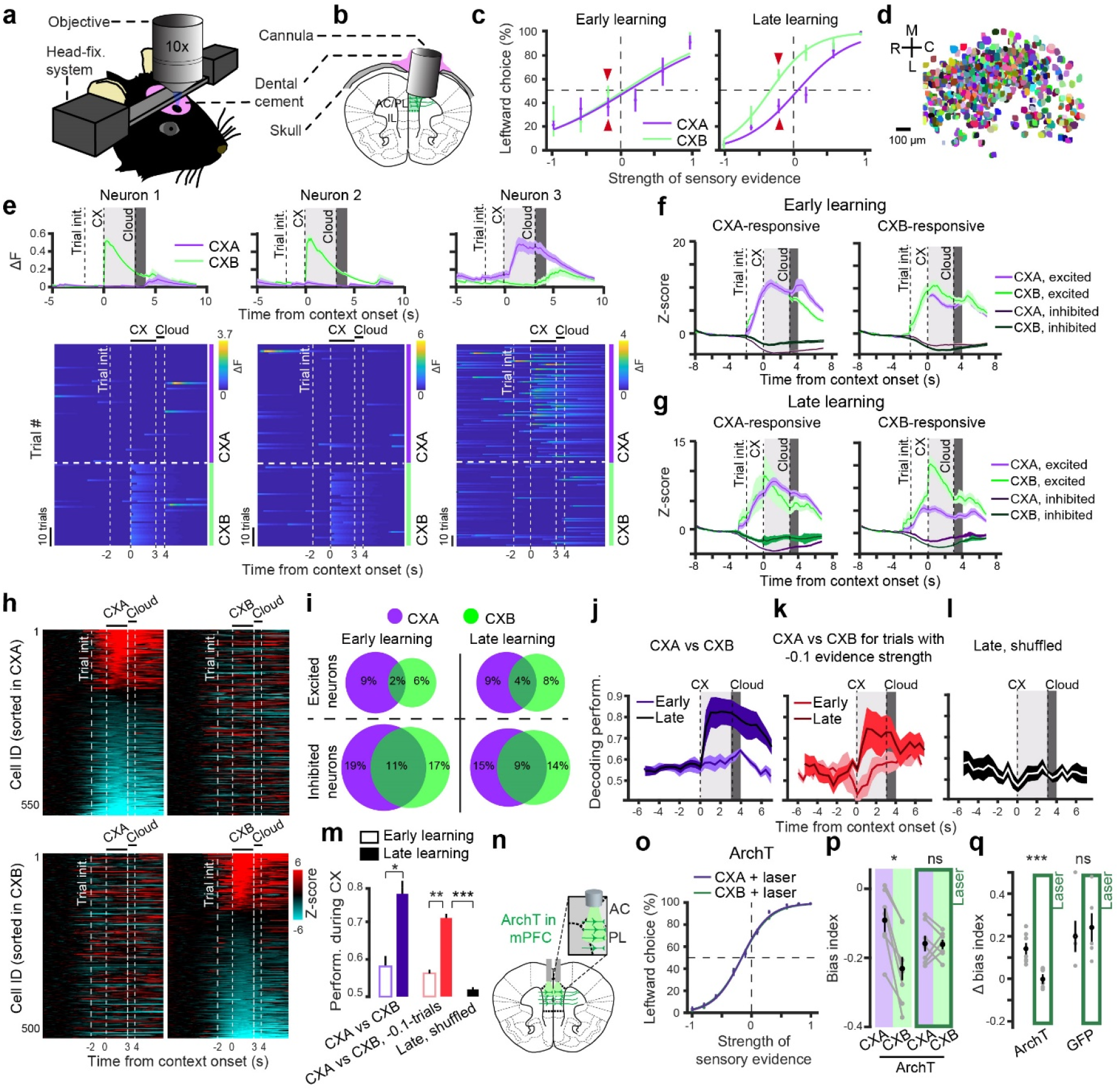
Learning-induced response in mPFC neurons essential for context-guided decisions. **(a, b)** Schematics of the imaging setup (a) and cannula implantation (b). **c,** Average psychometric curves for mice at the early (left, n = 4) and late (right, n = 6) training stages in the CX2AC task. **d,** Neurons detected and isolated from an example mouse. **e**, Example neurons responding to CXB (neuron 1 and 2) or CXA (neuron 3) at the late learning stage. Top panel, average activity over all trials in each context. Bottom panel, heatmaps of trial-by-trial activity in each context. **f,** Average activity of CXA-responsive (left) or CXB-responsive (right) neurons in different trial types. Activity was acquired during the early learning stage. **g,** Same as (f) but for activity acquired during the late learning stage. **h,** Heatmaps of the activity (z- scored) of the individual neurons in (g). Each row represents a neuron. Top: neurons sorted according to their activity during CXA. Bottom: neurons sorted according to their activity during CXB. **i,** Venn diagrams showing the percent distributions of neurons significantly (permutation test, P < 0.05) excited (top) or inhibited (bottom) by different contexts during early (left) and late (right) learning stages. Numbers indicate the overall percentages of neurons. **j,** SVM classifier performance for classifying CXA trials vs CXB trials during the early (n = 4 mice) and late (n = 6 mice) stages of learning. **k,** SVM classifier performance for classifying CXA trials versus CXB trials where the sensory evidence strength was the same (−0.1; indicated with red arrowheads in d) but mice made opposite choices, during the early (n = 4 mice) and late (n = 6 mice) stages of learning. **l,** Classification as in (k) but using trials shuffled between the two conditions. **m,** Quantification of performance during the contextual period for the results in (j-l) (*P = 0.0142, **P = 0.0069, ***P = 6.69e^−6^; paired t-test). **n,** Schematics of the approach for optogenetic inhibition of mPFC neurons. **o,** Psychometric curves in different contexts for mice (n = 6) in which the mPFC neurons were photo-inhibited by laser during context presentation. **p,** Quantification of bias indices in CXA and CXB for the ArchT mice in (o), at baseline (left) and when the mPFC neurons were photo-inhibited during the contextual period (right) (n = 7, *P = 0.0378), ns (nonsignificant), P = 0.901, paired t-test). **q,** Quantification of change in bias index caused by the contextual change. Photo-inhibition of mPFC neurons reduced the change (ArchT mice, n = 7, ***P = 0.00058; GFP mice, n = 4, ns, P = 0.7099; paired t-test). Psychometric curves are averaged over six sessions. Data are presented as mean ± s.e.m.

We simultaneously imaged the responses of large populations of neurons, for up to 655 neurons per mouse (381 ±190.69 (mean ±SD), n = 6 mice, for a total of 2286 neurons; Fig. 2d; Extended Data Fig. 1b; Extended Data Fig. 2a, b). At the late learning stage, we observed that individual neurons responded to either CXA or CXB (Fig. 2e), with the overall average response to CXB being higher than that to CXA (Extended Data Fig. 2c, d). This difference was not observed at the early training stage. We further analyzed the CXB- or CXA-responsive neurons, defined as the neurons showing significant response (excitatory or inhibitory; P < 0.05, permutation test) to presentations of CXB or CXA, respectively. Notably, at the late, but not early stage of training, the CXB-excited neurons displayed stronger excitatory response to CXB than CXA (Fig. 2f, g; Extended Data Fig. 2e-g). In contrast, the CXA-excited neurons displayed similar excitatory response to CXA and CXB at both stages (Fig. 2f, g; Extended Data Fig. 2e-g). It is noteworthy that this increased activity during CXB was consistent across trials with different cloud-of-tones stimuli (Extended Data Fig 2h, i). At both the early and late training stages, there were sizable CXB- or CXA-inhibited populations (Fig. 2f-i; Extended Data Fig. 2a, b, e-g) that responded differently to the two contexts (Fig. 2f, g; Extended Data Fig. 2e, f).

Since mPFC neurons showed a selective change in their response to CXB only after mice fully learned to adaptively switch between the newer, CXB-guided “go left” rule and the original, CXA-guided “follow the cloud” rule, these neurons may contribute to the context-guided rule selection. Indeed, decoding analysis with linear classifiers (support vector machine, SVM) revealed that mPFC neuron activity during the contextual period discriminated between CXA and CXB with high accuracy at a single trial level, and at the late but not at the early stage of training (Fig. 2j, m), suggesting that mPFC neurons encode the learned behavioral strategies. Similar results were obtained when the analysis was repeated on the trials in which the same cloud was presented and mice made opposite choices under the two contexts (Fig. 2k, l, m; trials marked with red arrows in c). These results indicate that the neuron activity predicts mice’s choices rather than tone-cloud categories. Interestingly, mPFC neurons also encoded the clouds specifically in the late learning phase (Extended Data Fig. 2j, k). Together, these results suggest that mPFC neurons encode the contexts and participate in context-guided decisions.

To determine if mPFC neurons are also required for context-guided decisions, we optogenetically inhibited these neurons during the contextual period in the CX2AC task (Fig. 2n; Extended Data Fig. 2l, m). We found that this manipulation completely abolished mice’s leftward bias under CXB (Fig. 2o-q). In contrast, optogenetic inhibition of mPFC neurons in the absence of the contextual cues had no effect on mice’s performance in the 2AC task (Extended Data Fig. 2n-p), suggesting that this manipulation does not alter general cognitive functions such as attention. Our results hence indicate that mPFC neurons not only participate in, but also are essential for context-guided strategy selection.

## mPFC^DMS^ neurons encode context-dependent decisions

The mPFC sends a vast network of top-down projections to many subcortical areas, among which the DMS has been shown to encode goal-directed behavior^22^. We reasoned that the DMS-projecting mPFC (mPFC^DMS^) neurons participate in the context-guided strategy selection. To test this hypothesis, we selectively targeted mPFC^DMS^ neurons for imaging with an intersectional viral strategy, by injecting the DMS with a retrograde AAV expressing Cre, and injecting the ipsilateral mPFC of the same mice with an AAV expressing GCaMP6 in a Cre-dependent manner (Fig. 3a; Extended Data Fig. 3a-c). These mice were implanted with cannulae in the mPFC and, after viral expression, were trained in the CX2AC task as described above (Fig. 3b; Extended Data Fig. 3d, e).

**Fig. 3.**
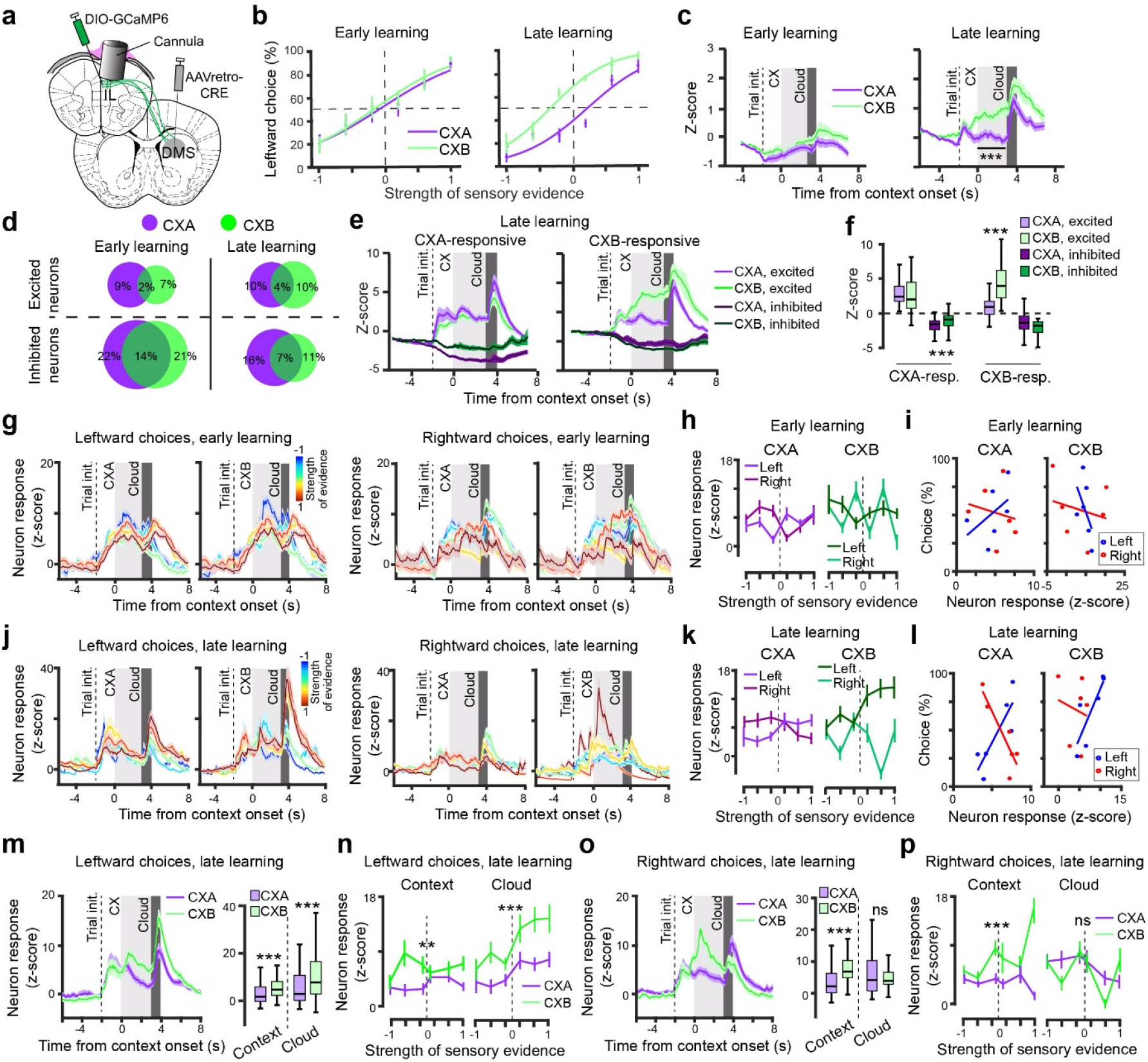
mPFC^DMS^ neuron activity represents context-dependent decisions. **a,** Schematics of the experimental design. **b,** Average psychometric curves for mice at the early (left, n = 5) and late (right, n = 6) training stages in the CX2AC task. **c,** Average activity of all neurons in CXA and CXB trials in the early (left) and late (right) learning stages (late, F_(1,30)_ = 68.677, ***P = 4.354e-15, two-way ANOVA with repeated measures). **d,** Venn diagrams showing the percent distributions of neurons significantly (permutation test, P < 0.05) excited (top) or inhibited (bottom) by different contexts during early (left) and late (right) learning stages. Numbers indicate the overall percentages of neurons. **e,** Average activity of CXA-responsive (left) or CXB-responsive (right) neurons in different trial types. Activity was acquired during the late learning stage. **f,** Quantification of the average activity during context presentation in (e) (CXA- responsive neurons: excitation, P = 0.62, inhibition, ***P = 5.9e-6, paired t-test; CXB- responsive neurons: excitation, ***P = 0.2.46e-6, inhibition, P = 0.08, Wilcoxon rank sum test). **(g-i)** Data acquired during the early learning stage. **g,** Activity of CXB-excited neurons in trials in which mice made leftward choices (left panel) and rightward choices (right panel). Each trace represents the averaged activity in trials with the same strength of sensory evidence. **h,** Responses of neurons in (g) to the cloud-of-tones stimuli as a function of evidence strength, separately plotted for different choices under either CXA (left) or CXB (right). **i,** Relationship between neuron responses to the cloud-of-tones stimuli (derived from h) and choices under either CXA (left) or CXB (right) (CXA: leftward, r = 0.468, P = 0.34, rightward, r = −0.19, P = 0.717; CXB: leftward, r = −0.434, P = 0.389, rightward, r = −0.254, P = 0.626; Pearson correlation analysis). **(j-p)** Data acquired during the late learning stage. **j, k** & **l,** same as g, h & i, respectively, except that data were acquired during the late learning stage. **l,** Relationship between neuron responses to the cloud-of-tones stimuli (derived from k) and choices under either CXA (left) or CXB (right) (CXA: leftward, r = 0.843, P = 0.034, rightward, r = −0.882, P = 0.0199; CXB: leftward, r = 0.936, P = 0.006, rightward, r = −0.266, P = 0.609; Pearson correlation analysis). **m,** Left panel: average activity of CXB-excited neurons in trials in which mice made leftward choices under CXA or CXB. Right panel: quantification of the activity during the context and cloud period (context, ***P = 3.07e-5, cloud, ***P = 2.6e-4, Wilcoxon rank sum test). **n,** Quantification of neuron activity during context and cloud periods as a function of evidence strength (context, F(1,88) = 8.5, **P = 0.004, cloud, F(1,88) = 12.3, ***P = 0.0005, Friedman test). **o & p,** Same as m & n, respectively, except that trials in which mice made rightward choices were analyzed. **o,** Context, *** P = 2.2e-8, cloud, P = 0.83, Wilcoxon rank sum test. **p,** Context, F(1,88) = 28.12, ***P = 2.8e-4, cloud, F(1,88) = 0.55, ns, P = 0.45, Friedman test. Data are presented as mean ± s.e.m. or box-and-whisker plots.

We imaged the activity of mPFC^DMS^ neurons during the CX2AC task at different stages of training (early stage, 696 neurons in 5 mice; late stage, 550 neurons in 6 mice; Fig. 3c; Extended Data Fig. 3f; Extended Data Fig. 4a). On average, these neurons showed sustained activation during CXB presentation at the late stage of training, which reached a peak following cloud presentation (Fig. 3c). Notably, the activity during CXB presentation was much higher than during CXA presentation at the late stage of training but not at the early stage (Fig. 3c; Extended Data Fig. 4a).

We sorted mPFC^DMS^ neurons into either CXB- or CXA-responsive (P < 0.05, permutation test) groups, and analyzed each group’s response to both contexts. Training significantly increased the percentage of CXB-excited neurons (p = 0.02, χ^2^ test,) but did not change the percentage of CXA-excited neurons (p = 0.74, χ^2^ test) (Fig. 3d). Interestingly, training generally decreased the percentage of both CXB-inhibited neurons (p = 6.01 × 10^−6^, χ^2^test) and CXA-inhibited neurons (p = 0.006, χ^2^ test) (Fig. 3d; Extended Data Fig. 4c-f).

At the late training stage, CXB-excited neurons clearly differentiated CXB from CXA, as they showed sustained and ramping-up activation during CXB presentation, up to much higher levels than their activity during CXA presentation (Fig. 3e, f; Extended Data Fig. 4d, f). These neurons did not discriminate between CXB and CXA at the early stage of training (Extended Data Fig. 4b, c, e). In contrast, CXA-excited neurons did not differentiate CXB from CXA at either late or early stage of training (Fig. 3e, f; Extended Data Fig. 4b-f). These results demonstrate that, through learning, mPFC^DMS^ neurons acquire ramping-up excitatory responses specific to CXB.

We reasoned that such CXB-specific response might instruct CXB-dependent decisions. To test this hypothesis and better understand how mPFC^DMS^ neuron response is related to mice’s decisions under different contexts, we identified all the neurons activated (z-score > 3) during the cloud period and analyzed the relationship between their activity and the animal’s choices under either CXA or CXB. In the early training stage, these neurons (n = 68) responded similarly to the clouds regardless of the choice or context (Fig. 3g, h), with the response not correlated with choice selection under either CXA or CXB (Fig. 3i). Strikingly, in the late training stage, these neurons (n = 88) displayed markedly different response profiles (Fig. 3j-l). Under CXA, their response to the clouds scaled with the evidence strength during either leftward or rightward choices (Fig. 3j, k). As a result, the amplitudes of responses to different clouds correlated with the probabilities of making leftward or rightward choices (Fig. 3l). However, under CXB, such predictive relationships were present only for leftward choices; for rightward choices, the cloud response of these neurons had no apparent relationship with either evidence strength or mice’s choices (Fig. 3j-l). These results suggest that the cloud response of PFC^DMS^ neurons in well-trained mice guides decision making in a context-dependent manner. Under CXA (and the “follow the cloud” rule), the response preserves instructive information for making both leftward and rightward choices; however, under CXB (and the “go left” rule), the information is present for leftward choices but absent for rightward choices.

To further understand how the contexts influence decision-related cloud response in PFC^DMS^ neurons, we analyzed these neurons’ activity during both the context period and the cloud period in different choices (Fig. 3m-p). For leftward choices in the late training stage, PFC^DMS^ neurons showed increased activity following CXB presentations, and further enhanced their activity upon cloud presentations. Consequently, the response to both the context and the clouds (including clouds with varying evidence strength) was markedly higher under CXB than under CXA (Fig. 3m, n; Extended Data Fig. 5c-h). In contrast, for rightward choices, although these neurons had increased activity following CXB presentations, their activity was reduced during the subsequent cloud presentations (Fig. 3o, p; Extended Data Fig. 5c-h). Of note, in the early training stage, the context-induced changes in PFC^DMS^ activity were either small or absent (Extended Data Fig. 5a, b).

Together, these results suggest that through learning, mPFC^DMS^ neurons acquire activity representing context-dependent rules, which can modulate decision making and influence choice behavior. In particular, CXB presentation, which instructed the “go left” rule, not only increased the activity of mPFC^DMS^ neurons during the contextual period, but also promoted their response to the clouds specifically during leftward choices (Fig. 3n). The latter effect is likely responsible for the leftward shift of the psychometric curve under CXB (Fig. 3b, late learning).

## mPFC^VMT^ neurons encode decisions independent of contexts

Since silencing the VMT (including the nucleus reuniens and adjacent nuclei) has been shown to impair contextual fear learning^12^ and strategy shifting^37^, we next tested whether VMT- projecting mPFC (mPFC^VMT^) neurons could also participate in context-instructed strategy switching. We targeted mPFC^VMT^ neurons using the intersectional viral strategy (Fig. 4a; Extended Data Fig. 6a-c) and imaged the activity of these neurons in mice in the CX2AC task at different stages of training (early stage, 714 neurons; late stage, 572 neurons; Fig. 4b; Extended Data Fig. 6d). On average, there was an increase in mPFC^VMT^ neuron activity in response to both CXA and CXB after mice learned the task, with the activity coupled with each of the task-related events: task initiation, context presentation, and cloud presentation (Fig. 4c; Extended Data Fig. 6e). However, unlike mPFC^DMS^ neurons, mPFC^VMT^ neurons as a whole did not show any significant difference in activity in CXB trials compared with CXA trials, at either early or late stage of training (Fig. 4c; Extended Data Fig. 7a-f). Training did not change the percentage of CXA- or CXB-responsive neurons either (CXA-excited, p = 0.5222; CXA- inhibited, p = 0.6772; CXB-excited, p = 0.7043; CXB-inhibited, p =0.6577; χ^2^ test; Fig. 4d). In addition, CXA- or CXB-excited neurons did not show any preference to either context throughout training (Fig. 4e, f; Extended Data Fig. 7a, b).

**Fig. 4.**
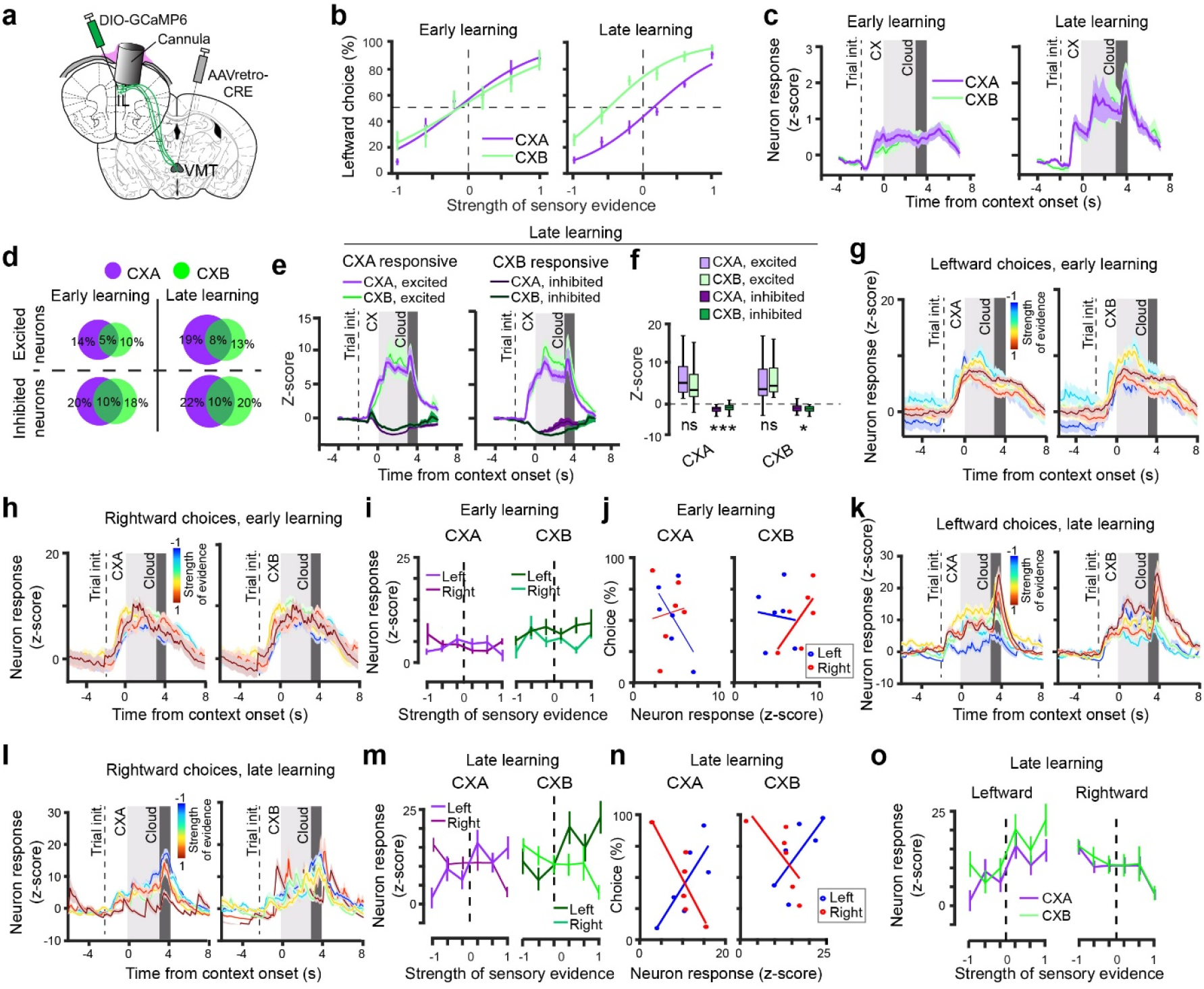
PFC^VMT^ neuron activity in the CX2AC task. **a,** Schematics of the experimental design. **b,** Average psychometric curves for mice at the early (left, n = 6) and late (right, n = 6) training stages in the CX2AC task. **c,** Average activity of all neurons in CXA and CXB trials in the early (left) and late (right) learning stages. **d,** Venn diagrams showing the percent distributions of neurons significantly (permutation test, P < 0.05) excited (top) or inhibited (bottom) by different contexts during early (left) and late (right) learning stages. Numbers indicate the overall percentages of neurons. **e,** Average activity of CXA-responsive (left) or CXB-responsive (right) neurons in different trial types. Activity was acquired during the late learning stage. **f,** Quantification of the average activity during context presentation in (e) (CXA-responsive neurons: excitation, P = 0.099, inhibition, ***P = 1.7e-3, paired t-test; CXB-responsive neurons: excitation, P = 0.167, inhibition, *P = 0.044, Wilcoxon rank sum test). **(g-j)** Data acquired during the early learning stage. **(g, h)** Activity of CXB-excited neurons in trials in which mice made leftward choices (g) and rightward choices (h). Each trace represents average activity in trials with the same strength of sensory evidence. **i,** Responses of neurons in (g, h) to the cloud- of-tones stimuli as a function of evidence strength, separately plotted for different choices under either CXA (left) or CXB (right). **j,** Relationship between neuron responses to the cloud-of- tones stimuli (derived from i) and choices under either CXA (left) or CXB (right) (CXA: leftward, r = −0.621, P = 0.188, rightward, r = 0.836, P = 0.109; CXB: leftward, r = 0.01, P = 0.97, rightward, r = 0.766, P = 0.075; Pearson correlation analysis). **(k-o)** Data acquired during the late learning stage. **k-n,** same as g-j, respectively, except that data were acquired during the late learning stage. **n,** Relationship between neuron responses to the cloud-of-tones stimuli (derived from m) and choices under either CXA (left) or CXB (right) (CXA: leftward, r = 0.796, P = 0.058, rightward, r = −0.85, P = 0.031; CXB: leftward, r = 0.819, P = 0.045, rightward, r= −0.856, P = 0.029; Pearson correlation analysis.) **o,** Quantification of neuron activity during cloud periods as a function of evidence strength, for both rightward choices and leftward choices (leftward, F(1,68) = 0.94, P = 0.31, rightward, F(1,68) = 1.77, P = 0.18, Friedman test). Data are presented as mean ± s.e.m. or box-and-whisker plots.

To determine if mPFC^VMT^ neuron response would be related to mice’s decision under different contexts, we identified all the neurons activated (z-score > 3) during the cloud period, and analyzed the relationship between their response and animal’s choice under either CXA or CXB. Like mPFC^DMS^ neurons, in the early training stage, PFC^VMT^ neurons responded similarly to the clouds regardless of the choice or context (Fig. 4g-i), and the response did not correlate with choice selection under either context (88 neurons; Fig. 4j). However, in the late training stage, the cloud response of mPFC^VMT^ neurons (n = 111) became predictive of the tones in the clouds (Fig. 4k-m), and of animal’ s choices (Fig. 4n), under both CXA and CXB. In particular, under CXB, the activity of these neurons correlated with not only leftward choices, but also rightward choices (Fig. 4m, n). In fact, the cloud–response relationship during either the rightward or the leftward choices were highly similar between CXB and CXA conditions (Fig. 4o). This observation is markedly different from that of PFC^DMS^ neurons, which did not show such predictive properties during rightward choices under CXB (Fig. 3l, p).

A fraction of mPFC^VMT^ neurons was inhibited by context or cloud presentations across training (Fig. 4d-f; Extended Data Fig. 7; Extended Data Fig. 8a-d). Interestingly, we found that the inhibitory response during leftward choices was more potent under CXB than under CXA, specifically in the late phase of training (Extended Data Fig. 8a-d). Such difference was absent during rightward choices (Extended Data Fig 8b, d), and was also not observed in mPFC^DMS^ neurons throughout training (Extended Data Fig. 8e-h).

Together, these results suggest that mPFC^VMT^ neurons have very different encoding properties compared with mPFC^DMS^ neurons. In particular, a subset of PFC^VMT^ neurons show learning-dependent excitatory response that does not represent contextual changes and the associated rule switch. Instead, these neurons represent the old “follow the cloud” rule even though mice make choices according to the new “go left” rule under CXB. In other words, these neurons seem to keep a fixed representation of the original strategy and its associated choice despite the actual changes in strategy and choice. In contrast, another subset of mPFC^VMT^ neurons show learning-dependent increase in inhibition following CXB presentation specifically during left choices, suggesting that inhibition of these neurons is important for the CXB-dependent change in strategy.

## mPFC^DMS^ and mPFC^VMT^ have opposing roles in context-dependent decisions

Since mPFC^DMS^ neuron activation following CXB presentation predicts the switch of behavioral strategy from “follow the cloud” to “go left”, we reasoned that suppressing this neuronal activation would impair the strategy switch. On the other hand, mPFC^VMT^ neuron activation stably represents the original strategy, but inhibition of a subpopulation of mPFC^VMT^ neurons is associated with the strategy switch. Hence, it is possible that inhibition of mPFC^VMT^ neurons might facilitate the switch.

To test these predictions, we used optogenetics based on *Guillardia theta* anion-conducting channelrhodopsins (GtACR)^52,53^ to inhibit mPFC^DMS^ or mPFC^VMT^ neurons (Fig. 5a-d; Extended Data Fig. 9a, b). We found that inhibition of mPFC^DMS^ neurons specifically during the contextual period in the CX2AC task caused a reduction in leftward choice under CXB, especially in the trials in which mice were most influenced by the “go left” rule (i.e., the trials with clouds of −0.1 evidence strength; Fig. 5b). The same manipulation did not affect behavior once the contextual cues were omitted (Extended Data Fig. 9c). In sharp contrast, inhibition of mPFC^VMT^ neurons during the contextual period led to an increase in leftward choice under CXB, specifically in the trials with clouds of −0.1 evidence strength (Fig. 5d). This effect disappeared when the contextual cues were absent (Extended Data Fig. 9d). Of note, inhibiting mPFC^DMS^ or mPFC^VMT^ neurons had no effect on CXA trials (Fig. 5b, d). Control experiments showed that light illumination in the mPFC alone had no effect on behavior in the CX2AC task (Fig. 5e-h; Extended Data Fig. 9e, f). Thus, inhibiting mPFC^DMS^ or mPFC^VMT^ neurons during the presentation of CXB prevents or facilitates, respectively, the use of the CXB-guided “go left” rule. These results suggest that mPFC neurons are causally involved in context-dependent selection of strategy, with mPFC^DMS^ neurons being critical for the selection of a newly learned rule whereas mPFC^VMT^ neurons favoring the use of the original rule.

**Fig. 5.**
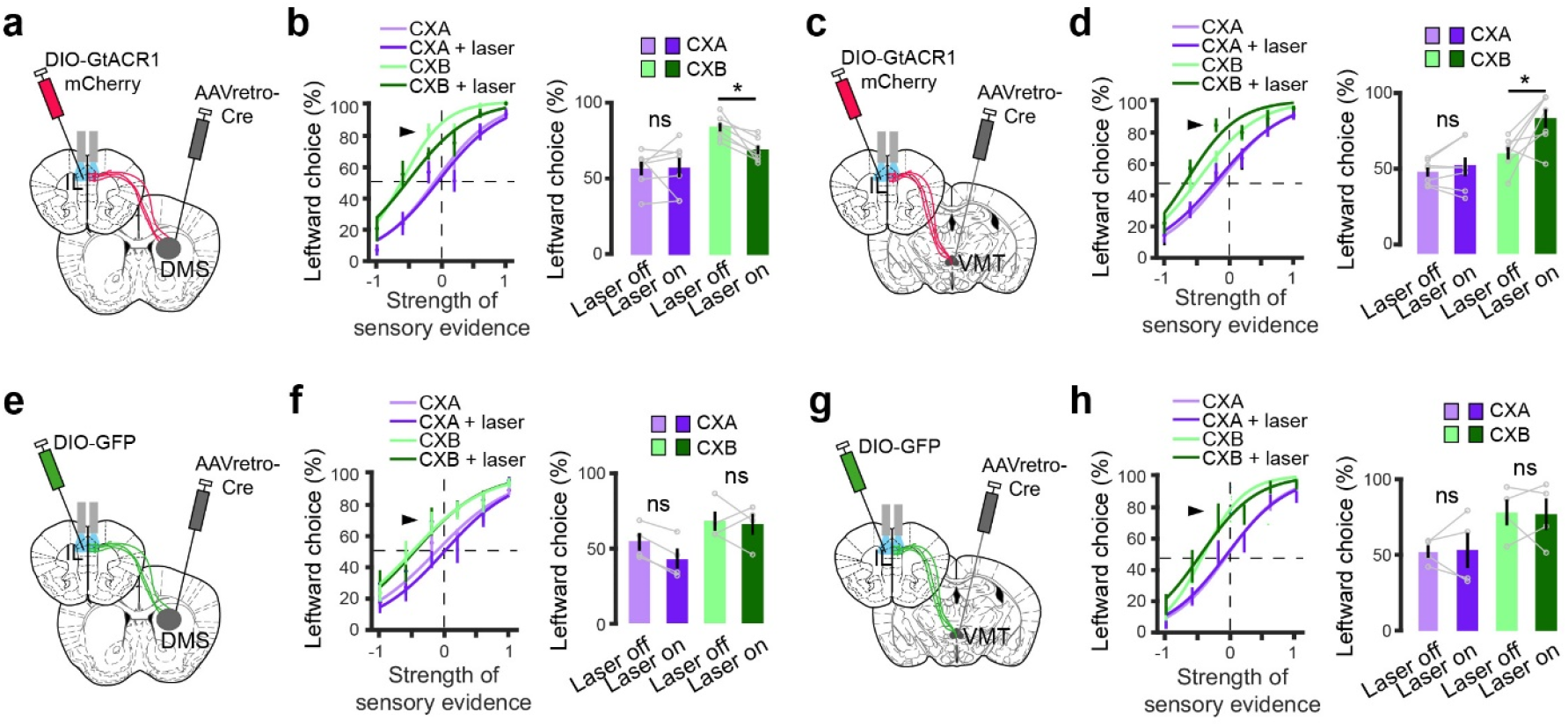
Inhibiting mPFC^DMS^ or mPFC^VMT^ oppositely influences context-dependent decisions. **a,** Schematic of the experimental design. mPFC^DMS^ neurons were optogenetically inhibited during context presentation in the CX2AC task. **b,** Left panel: psychometric curves of the GtACR1 mice for different trial types. Data represent the average of 3 sessions from 7 mice. Right panel: quantification of leftward choices in trials where the sensory evidence strength was −0.1 (indicated by the arrowhead on the psychometric curves) (F_(1,6)_ = 18.54, P = 0.001; ns (nonsignificant), P > 0.05; *P < 0.05; two-way ANOVA with repeated measures, followed by Student paired t-tests with Bonferroni corrections). **c,** Schematic of the experimental design. mPFC^VMT^ neurons were optogenetically inhibited during context presentation in the CX2AC task. **d,** Left panel: psychometric curves of the GtACR1 mice for different trial types. Data represent the average of 3 sessions from 7 mice. Right panel: quantification of leftward choices in trials where the sensory evidence strength was −0.1 (indicated by the arrowhead on the psychometric curves) (F(1,6) = 16.01, P = 0.0018; ns, P > 0.05; *P < 0.05; two-way ANOVA with repeated measures, followed by Student paired t-tests with Bonferroni corrections). (**e, f**) Same as (a, b), except that GFP was expressed in mPFC^DMS^ neurons. **f,** Left panel: psychometric curves of the GFP mice for different trial types. Data represent the average of 3 sessions from 4 mice. Right panel: quantification of leftward choices in trials where the sensory evidence strength was −0.1 (indicated by the arrowhead on the psychometric curves) (F(1,3) = 10.7, P = 0.017; ns, P > 0.05; two-way ANOVA with repeated measures, followed by Student paired t-tests with Bonferroni corrections). (**g, h**) Same as (c, d), except that GFP was expressed in mPFC^VMT^ neurons. **h,** Left panel: psychometric curves of the GFP mice for different trial types. Data represent the average of 3 sessions from 4 mice. Right panel: quantification of leftward choices in trials where the sensory evidence strength was −0.1 (indicated by the arrowhead on the psychometric curves) (F(1,3) = 8.7, P = 0.0256; ns, P > 0.05; two-way ANOVA with repeated measures, followed by Student paired t-tests with Bonferroni corrections). Data are presented as mean ± s.e.m.

## Discussion

In the CX2AC task, mice learned that one context (CXA) is associated with a previously learned rule (the “old rule”) that requires decisions be solely based on sensory evidence, while another context (CXB) is associated with a newly learned rule, which requires that decisions be made on the basis of both the sensory evidence and reward values. Thus, this task captured critical features of adaptations that animals and humans make in real life, including the adjustment of decisions according to reward values, the learning of a new rule on top of an already established one, and the use of contexts to guide strategy selection.

The CX2AC task combined with *in vivo* imaging and optogenetics allowed us to demonstrate, to our knowledge, for the first time that individual mPFC neurons acquire response through learning that encodes contexts and is necessary for context-dependent selection of decision strategies. The activity of a subset of mPFC^DMS^ neurons discriminated sensory stimuli and predicted choices when mice make decisions based on sensory evidence. Importantly, these neurons showed increased response to CXB compared with that to CXA, and further adapted their response during subsequent sensory discrimination under CXB, with the response increasing to higher levels and predicting choices if mice selected the correct decision strategy. Moreover, optogenetic inhibition of mPFC^DMS^ neurons during CXB presentation prevented the strategy selection. These results suggest that a brief exposure to contextual cues profoundly modulates the subsequent response profile of mPFC^DMS^ neurons during decision making, thereby influencing choices.

In contrast, although a subset of mPFC^VMT^ neurons increase context-related activity with learning, these neurons do not discriminate between CXA and CXB. In addition, these neurons have similar activity profiles during sensory discrimination under the old and new rules, responding as if the old rule were in place and the original decision strategy were followed, even when the rule has changed and the mice have adopted the new strategy. Interestingly, the subset of mPFC^VMT^ neurons showing inhibitory response does discriminate between contexts, with the inhibition being more potent during CXB than CXA. Furthermore, optogenetic inhibition of mPFC^VMT^ neurons during CXB facilitated switching to the new strategy. These results suggest that a decrease in activity in mPFC^VMT^ neurons may permit switching to the new rule.

Altogether, our results suggest that the two mPFC top-down projections exert distinct and complementary functions in decision making: while mPFC→DMS projections support flexible context-guided strategy selection, mPFC→VMT projections keep the stability of decision strategies. A balance of activity between the two pathways may be critical for adaptive behaviors.

## Acknowledgements

We thank Christian Bravo-Rivera, Walter Bast, Jonathan Cahn, and Benoît von der Weid for comments on an earlier version of the manuscript, and members of the Li laboratory for helpful discussions. This work was supported by grants from NARSAD (26276, to O.G.; 28229, to X.Z.), Swiss National Science Foundation (P300PB-174497, to O.G.), the National Institutes of Health (NIH) (R01MH101214, R01MH108924, R01NS104944, R01DA050374, to B.L.), Human Frontier Science Program (RGP0015/2016, to B.L.), the Cold Spring Harbor Laboratory and Northwell Health Affiliation (to B.L.), and Feil Family Neuroscience Endowment (to B.L.).

## Author contributions

O.G. and B.L. conceived and designed the study. O.G. conducted the experiments and analyzed data. D.V.D.L., T.Y., X.Z. and R.S. assisted with experiments and analysis. T.Y. developed the one-photon wide-field imaging system and methods. O.G. and B.L. wrote the paper with inputs from all authors.

## Material and Methods

### Animals

Male and female mice with age of 2-4 months were used in all the experiments. Mice were housed under a 12-h light/dark cycle (7 a.m. to 7 p.m. light) in groups of 2-5 animals, with the room temperature being 22°C and humidity being 50%. Food and water were available *ad libitum* before behavioral training. All behavioral experiments were performed during the light cycle. Littermates were randomly assigned to different groups prior to experiments. All mice were C57BL/6J. All experimental procedures were approved by the Institutional Animal Care and Use Committee of Cold Spring Harbor Laboratory (CSHL) and performed in accordance to the US National Institutes of Health guidelines.

### Stereotaxic surgery

All surgery was performed under aseptic conditions and body temperature was maintained with a heating pad. Standard surgical procedures were used for stereotaxic injection and implantation as previously described^50,54^. Briefly, mice were anesthetized with isoflurane (2% in a mixture with oxygen, applied at 1.0 L/min), and head-fixed in a stereotaxic injection frame, which was linked to a digital mouse brain atlas to guide the targeting of different brain structures (Angle Two Stereotaxic System, myNeuroLab.com). Lidocaine was injected subcutaneously in the head and neck area as a local anesthetic.

We first made a small cranial window (1-2 mm^2^) above the target brain region. For imaging or optogenetic inhibition of mPFC neurons, we lowered a glass micropipette (tip diameter, ~5 μm) containing the AAV1.Syn.GCaMP6f.WPRE.SV40 or rAAV9/CAG-ArchT-GFP viral solution, respectively, to reach the mPFC (coordinates: 1.8 mm anterior to Bregma, 0.3 mm lateral from midline, and 2.3 mm vertical from brain surface). About 0.3-0.4 μl of viral solution was delivered with pressure applications (5-20 psi, 5-20 ms at 1 Hz) controlled by a Picospritzer III (General Valve) and a pulse generator (Agilent). The rate of injection was ~20 nl/min. The pipette was left in place for 10 min following the injection, and then slowly withdrawn.

To target the PFC^VMT^ pathway for imaging and optogenetics, we injected a retrograde AAV (rAAV2-retro-Syn-Cre (HHMI-Janelia Research Campus) for imaging, or pAAV-Ef1a- mCherry-IRES-Cre (Addgene) for optogenetics; 0.3-0.4 μl) bilaterally into two different locations of the VMT (coordinates: (1) 1.46 mm posterior to Bregma, 0.0 mm lateral from midline, and 3.85 mm vertical from brain surface; (2) 0.82 mm posterior to Bregma, 0.0 mm lateral from midline, and 3.8 mm vertical from brain surface). To target the PFC^DMS^ pathway for imaging and optogenetics, we injected the same viruses (0.4-0.5 μl) bilaterally into two different locations of the DMS (coordinates: (1) 0.86 mm anterior to Bregma, 1.3 mm lateral from midline, and 3.8 mm vertical from brain surface; (2) 0.38 mm anterior to Bregma, 1.4 mm lateral from midline, and 3.85 mm vertical from brain surface). We then injected the pAAV.Syn.Flex.GCaMP6f.WPRE.SV40 (Addgene; 0.3-0.4 μl; unilateral for imaging), or AAV_hSyn1-SIO-stGtACR1-FusionRed (Addgene; 0.3-0.4 μl; bilateral for inhibition) into the mPFC (coordinates: 1.8 mm anterior to Bregma, 0.3 mm lateral from midline, and 2.3 mm vertical from brain surface).

For optogenetics, we further implanted optic fibers bilaterally 200-300 μm above the injection locations in the mPFC with a 6° angle using an optic fiber holder (ThorLabs). The optic fibers (core diameter, 200 μm; length, 3 or 4 mm; NA, 0.22; Inper, Hangzhou, China) used for the photostimulation transmitted light with >90% efficiency when tested before implantation. Optic fibers were attached to the skull using a UV light-sensitive dental cement (3M RelyX Unicem). A home-made stainless-steel head-bar was also mounted next to the posterior part of the optic fiber for head restraint. Additional dental cement was added to seal the preparation.

To prepare mice for imaging, 3-5 days following the virus injection, the mice underwent another surgery as described above, during which we implanted a cannula with a glass coverslip at its bottom (two types of cannulae were used: (1) outer diameter, 1.8 mm; length, 4 mm; (2) outer diameter, 1 mm; length, 3 mm; Inscopix) into the mPFC. Before implanting the cannula, drops of diluted anti-inflammatory solution (dexamethasone, Metacan) were added to the cranial window and washed 30-60 s later with a saline solution. The cannula was slowly (~20 μm/min) lowered to the mPFC with a cannula holder, to depths that were above the viral injection location (coordinates: 1.8 mm anterior to Bregma, 0.3 mm lateral from midline, and 2.3-2.4 mm vertical from brain surface). The cannula was attached to the skull using UV light-sensitive dental cement. A head-bar (for head restraint) was subsequently mounted to the skull as described above. The skin of mice was sutured using medical glue (3M Vetbond Tissue Adhesive). We waited for at least 4 weeks before starting the imaging experiments in these mice.

### Behavioral apparatus

Mice were head-restrained with the head-bar on a home-made head-fixation system. Three metal spouts were placed approximately 5 mm below mouse’s mouth. The distance between the adjacent spouts was ~4 mm. The spouts were arranged such that the mice could reach each spout with their tongue. The spouts were made of needles (CML supply, industrial dispensing tips, 16 gauge, 1-1/2’’ long) connected to silicon tubes, which were further connected to 50-ml syringes containing water. Gravity flow of water through the tubes was controlled by electronic valves (Lee Company, LHD series solenoid valve). The spouts were held together using a 3D printed plastic holder^43^, which was attached to a 3-axis manual micromanipulator (Thorlabs, DT12XYZ). The placement of the spouts was adjusted with the micromanipulator, and was monitored with a webcam placed under the spouts. Each spout also served as part of a custom “lickometer” circuit, which registered a lick event each time a mouse completed the circuit by licking the spout.

Stimulus playback and trial control were performed via a Bpod/PulsePal open-source Arduino-based system (Sanworks, Stony Brook, NY, USA). Custom scripts written in MATLAB based on Bpod commands were used to control the delivery of different stimuli and record licking events. Auditory stimuli were uploaded to the audio adaptor board using the Bpod control system.

### Behavioral training

Mice were kept on a water-restriction schedule (1 ml of water per day for each mouse), starting 48 h before the onset of training in the 2AC task. The training protocol for the 2AC task was derived from previous studies^42,43,55^. A white light signaled the start of a trial. Mice were taught to initiate the trial sequence by licking the central spout, which was rewarded with 0.5 μl of water. Five seconds after the trial initiation, a 1-s “cloud-of-tones” stimulus was presented. Mice were trained to lick the left spout when hearing a cloud containing high-frequency tones (12-17kHz) and the right spout when hearing a cloud containing low-frequency tones (1-6kHz). A correct choice was rewarded by a 3-μl drop of water, whereas an error choice led to nothing in that trial. A choice was defined as correct if mice committed to licking the same spout during the second half of the cloud period. The white light was switched off when mice responded to the cloud, irrespective of the choice being correct or incorrect. The inter-trial interval (ITI) lasted for 10-15 seconds.

The tones in the cloud were drawn according to a protocol modified based on previously described ones^42,43,55^. The cloud consisted of a stream of 30-ms pure tones presented at a rate of 100 tones per second. We used eighteen possible tones with frequencies logarithmically spaced between 1 kHz and 18 kHz. On easy trials, tones were drawn exclusively from those with the target frequencies: low (1-6 kHz) for rightward choice, and high (12-18 kHz) for leftward choice. On intermediate trials, the clouds contained a mixture of tones with different frequency ranges. The difference in the rates between high-frequency tones and low-frequency tones in a cloud determined the strength of sensory evidence:
Strength of evidence for leftward choice = [Tones_high_ – Tones_low_] (tones per s) / 100 (total tones per s)

For example, a cloud with an evidence strength of 0.6 meant that for each time slot in the cloud, the chance for the tone to be picked from the high-frequency range (12-18 kHz) was 60% higher than the chance for it to be picked from the low-frequency range (1-6 kHz). We used clouds with evidence-strength values of −1, −0.6, −0.1, 0.1, 0.6, and 1 for all the imaging and pathway-specific optogenetics experiments. The negative values corresponding to the strengths favoring rightward choice.

We started by training mice with easy trials. The different types (leftward-choice and rightward-choice) of these trials were either randomly interleaved, or alternated between two different blocks, with each block containing 5 trials of the same type. Once mice reached a reasonable performance (70-80% correct trials, 5-20 days of training), intermediate trials with clouds of fixed evidence strength were gradually introduced. For imaging experiments, all mice were first trained in alternating blocks, with each block containing 5 trials of the same type (leftward-choice or rightward-choice). Mice were able to categorize the clouds at 3 weeks after training started and increased the performances over a 7-10-week training period.

After mice were able to discriminate the intermediate clouds, we started to train them in the CX2AC task by adding two distinct contextual cues to the 2AC task, context A (CXA) and context B (CXB). CXA consisted of a 3-s UV light and a 1-s 4.5-kHz pure tone. CXB consisted of a 3-s green light and a 1-s 12kHz pure tone. In each trial, one of the two contextual cues were presented at 2 s following the onset of trial initiation to indicate that one of two rules would be applied. CXA informed that the original rule (the “old rule”) would be in effect, so that the mice should keep making choices based on the sensory evidence in the subsequent cloud. CXB informed that a large reward (10 μl) would be delivered if mice made a correct leftward choice according to the old rule; however, no rightward choice would lead to any reward. Thus, under this “new rule”, mice still need to use the sensory evidence in the clouds in order to make correct leftward choices, but should ignore the clouds indicating a rightward choice under the old rule. We randomly interleaved CXA trials and CXB trials, but kept the former as the majority of trials (70% for all the mice in Fig. 1, 2 & 5, and 4 mice in Fig. 3 & 4; 60% for 4 mice in Fig. 3 & 4). After 3-6 weeks of training with the two contextual cues, ~80% of mice showed a substantial leftward bias in CXB trials but not in CXA trials in psychometric function. The bias was assessed by averaging the behavioral performance in 6 sessions for each mouse.

### Behavior analysis

The psychometric curve was generated by fitting a sigmoidal function using a built-in Matlab routine. The context-dependent bias index was computed as the strength of sensory evidence in the cloud (between −1 and 1) where the psychometric curve crossed the chance level, that is, where the fraction of leftward choice was equal to 50%. The change in bias index (Δ bias) was calculated as the bias index in CXB trials minus that in CXA trials. The fraction of leftward choice was calculated as the fraction in the psychometric curve corresponding to a strength of sensory evidence of 0.

### Optogenetic experiment

We used a green laser (532 nm, OEM Laser Systems Inc., Bluffdale, Utah, USA) for photoinhibition of mPFC neurons. We delivered the light bilaterally into the mPFC during the contextual period in all trials (power, 10-15 mW measured at the tip of optic fibers; 3-s constant light illumination between contextual cue onset and cloud onset). Data from six sessions were used for analysis.

For the pathway-specific photoinhibition, we delivered the light bilaterally into the mPFC during the contextual period in 30% of trials. We used data from three sessions that met the following criteria for analysis: (1) the bias index (see above) in CXA trials was not below - 0.3, indicating that mice did not have a substantial baseline bias towards the left side; and (2) the difference in the bias index between CXA trials and CXB trials was larger than 0.25, indicating that the mice used the contextual cues to guide decisions. Data were averaged across the three sessions for each mouse.

### Calcium imaging acquisition and analysis

All imaging experiments were conducted on behaving head-restrained mice in a dark, sound attenuated box. A custom-built wide-field imaging system was used to image GCaMP6 fluorescence signals. The system consisted of four major components: excitation light source, imaging optics, CCD camera and acquisition software, and mechanical parts. An LED (470 nm; PE-100, CoolLED) was used as the excitation light source. During imaging, the light power was adjusted to 0.1–0.4 mW, and was set to be constant for the same animal across imaging sessions. A filter cube (U-MF2, Olympus), which contained the appropriate optical filters, was used to ensure that only fluorescence signals with the desired wavelengths are transmitted. The filters were: excitation (FF02-482/18-25, Semrock), dichroic (FF409/493/573/652-Di01, Semrock) and emission (FF01-520/35-25, Semrock). An objective lens (10x, NA 0.3, WD 11 mm; MPLFLN10X, Olympus) was used to focus the excitation light, and collect fluorescence signals through the implanted cannulae. A tube lens (180 mm; TTL180-A, Thorlabs) was paired with the objective for magnification and forming images onto a monochrome CCD camera (pco, digital 14 bit CCD camera, image sensor ICX285AL, pco.pixelfly), which was used to collect fluorescence signals. A custom Imaging Acquisition software written in LabVIEW (National Instruments) was used to interface the camera with a dedicated desktop computer and record the GCaMP6 signals at a frame rate of 10 frames/s. To synchronize imaging with behavioral events, Imaging Acquisition was triggered with a TTL (transistor-transistor logic) signal from the Bpod State Machine (Sanworks) used for behavioral control. During imaging, the timestamps of different events, including the trigger signals sent to Imaging Acquisition, the onset of different stimuli, and licking events were all recorded with Bpod.

For imaging data processing and analysis, we first used Inscopix Data Processing software (v.1.2.0., Inscopix) to spatially down-sample all the raw images by a factor of 4 to reduce file size, and to correct the image stack for motion artifacts. The motion-corrected images were cropped to remove post-registration borders and margin areas. The pre-processed image stack was exported as a .tif file. Next, we used the extended constrained non-negative matrix factorization optimized for one-photon imaging (CNMF-E) ^50,51,54,56,57^ to demix neural signals and get their denoised and deconvolved temporal activity, termed ΔF^56,57^, for further analysis.

To determine whether a neuron was significantly (P < 0.05) excited or suppressed by a stimulus, and thus can be classified as being “responsive” to the stimulus, we used a permutation test to compare the mean ΔF values during baseline (the 3-s period immediately before trial initiation) with those during the contextual period (between 2 s and 5 s after trial initiation), or with those during both the contextual period and the cloud period (between 2 s and 6 s after trial initiation), taking all the trials into consideration. We used the Chi-square test to compare the percentages of responsive neurons before and after learning. For further analyses, we used z-scores to represent the dynamic activities in each neuron. To obtain the temporal z-scores for a neuron, we first obtained the mean activity trace for the neuron by averaging the fluorescence signals (ΔF) at each time point across all trials, and then computed the z-scores as (F(t)–F_mean_)/F_SD_, where F(t) is the ΔF value at time t, F_mean_, and F_SD_ are the mean and standard deviation, respectively, of the ΔF values over the baseline period. To analyze the responses of neuronal populations during leftward or rightward choices in trials with different cloud-of-tones stimuli, we selected individual neurons with z-scores higher than 3 during CXA or CXB period. We further averaged the responses of the selected neurons for each condition.

### Decoding analysis

We performed population decoding analysis using the linear support vector machine (SVM) in MATLAB (fitcsvm) to determine whether the types of trials could be predicted on the basis of the trial-by-trial population activities. We used the activities of all the simultaneously imaged neurons in each session in each mouse to perform the population decoding analysis. We used a subset of the low dimensional trial-by-trial neuronal activity data as the training dataset to train a classifier with linear kernel function (‘linear’) for two-class decoding (e.g., classifying CXA trials and CXB trials). Finally, we validated the classifier by using the ‘predict’ function to classify the trial-by-trial neuronal activities in the test dataset. Activities from randomly selected 80% of trials of each type were used to train the classifier, and activities from the remaining 20% of trials of each type were used to test decoding accuracy. To generate the shuffled data, we randomly reassigned a trial type to each of the trial-by-trial neuronal activities. We then followed the same procedure as that used for classifying the actual data to decode the shuffled data. We repeated this classification process 50 times for both the actual test dataset and the shuffled data, and calculated the average accuracy as the decoding accuracy.

### Statistical analysis

Statistical analyses were conducted using MATLAB statistical toolbox (MathWorks). The statistical test used for each comparison is indicated when used. Parametric tests were used whenever possible to test differences between two or more means. Non-parametric tests were used when data distributions were non-normal. Analysis of variance (ANOVA) was used to check for main effects and interactions in experiments with repeated measures, and for one or more factors. All comparisons were two tailed. Statistical hypothesis testing was conducted at a significance level of 0.05, with Bonferroni corrections when multiple tests were performed

## FIGURES, EXTENDED DATA FIGURES, and LEGENDS

**Extended Data Fig. 1.**
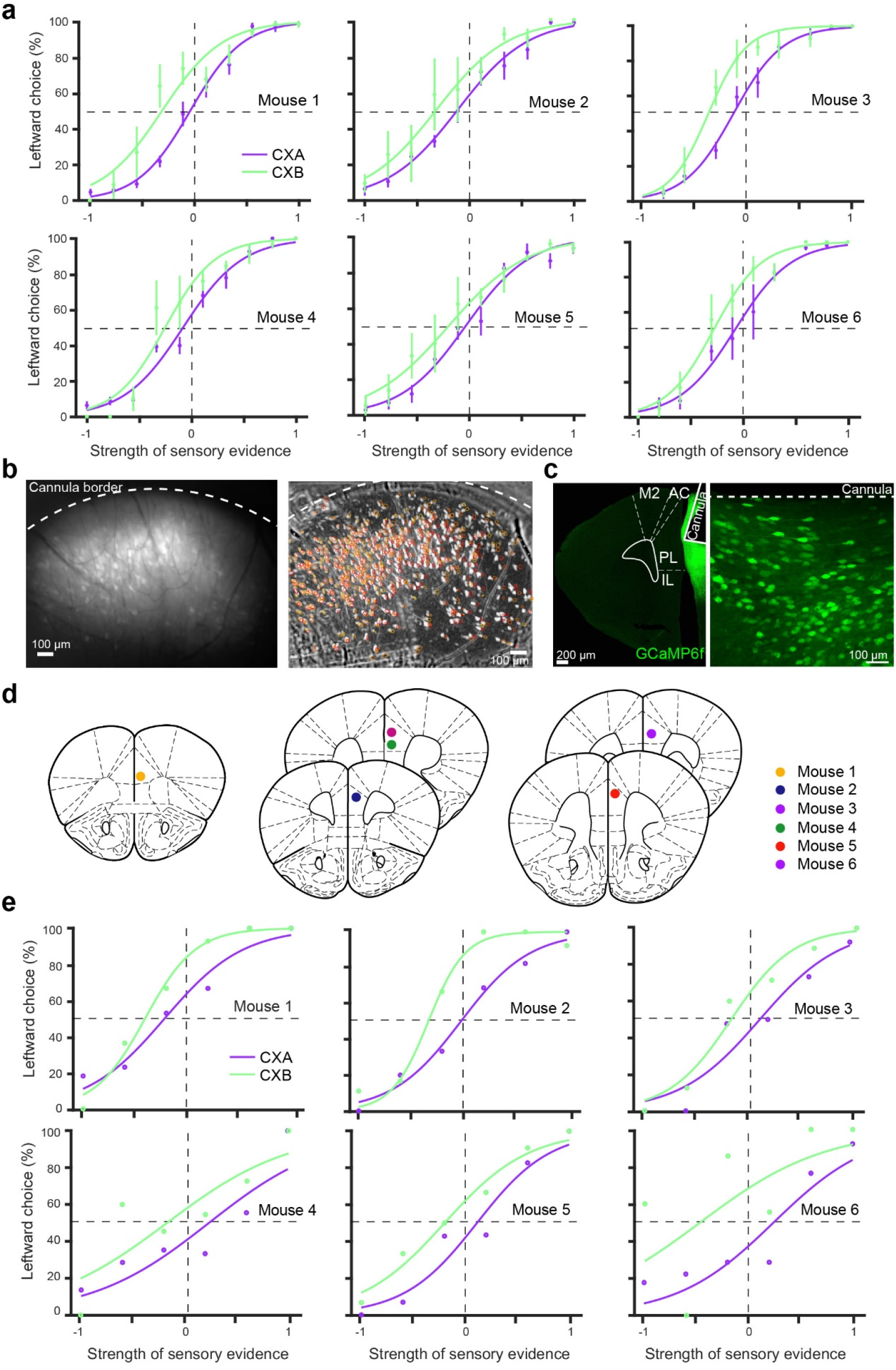
Behavioral performance and histology of the mice used for imaging mPFC neurons. a, Psychometric curves of individual mice. Performance of each mouse under different contexts was averaged over six sessions for each of the eight cloud-of-tones stimuli, which had evidence strength between −1 and 1 for the leftward choice. **b,** Images of the mPFC of an example mouse. Left: maximum intensity projection of raw images recorded during an imaging session. Dashed line represents the border of the cannula. Right: same field of view as that in the left, with isolated individual neurons marked with circles. **c,** Confocal images showing mPFC neurons expressing GCaMP6f and canula placement. On the right is a higher magnification image of the mPFC area on the left. **d,** Diagrams showing cannula placement in the mPFC of different mice. **e,** Psychometric curves of individual mice during imaging.

**Extended Data Fig. 2.**
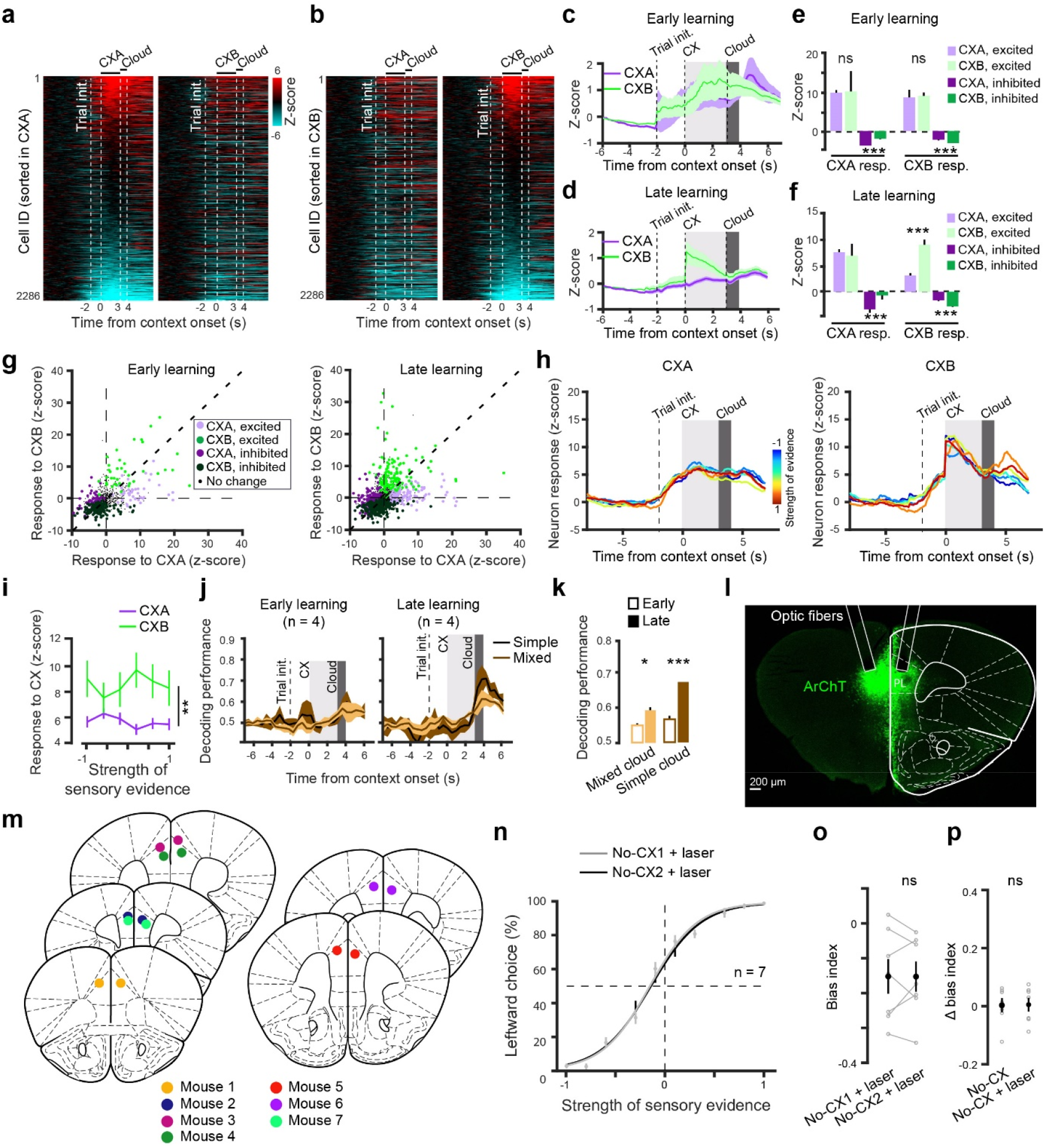
mPFC neurons encode contextual information in a learning-dependent manner. (**a, b**) Heatmaps of the activity (z-scored) of all individual neurons under different contexts (n = 2286 neurons, 6 mice). Each row represents a neuron. **a,** Neurons are sorted according to their activity during CXA. **b,** Same data as that in (a), but neurons are sorted according to their activity during CXB. (**c, d**) Average activity of all neurons in CXA and CXB trials in the early (c) and late (d) learning stages. **e,** Quantification of the average activity during context presentation in (c) (CXA-responsive neurons: excitation, ns (nonsignificant), P = 0.623, inhibition, ***P = 5.751e-10; CXB-responsive neurons: excitation, ns, P = 0.31, inhibition, ***P = 3.535e-6; Student paired t test). **f,** Quantification of the average activity during context presentation in (d) (CXA-responsive neurons: excitation, ns, P = 0.055, inhibition, ***P = 5.75e-11; CXB-responsive neurons: excitation, ***P = 1.0189e-7, inhibition, ***P = 8.6e-12; Student paired t test). **g,** Relationship between CXA-response and CXB-response for individual neurons displaying a significant activity change or no change (P < 0.05 or P > 0.05, respectively, permutation test) during the contextual period, in the early (left) and late (right) stages of training. **h,** Activity of CXA-excited neurons (left) and CXB-excited neurons (right) at the late stage of training, with each trace representing the average activity in trials with the same strength of sensory evidence. **i,** Quantification of neuron activity during the context period as a function of evidence strength, for both CXA trials and CXB trials (F(1,313) = 8.1, **P = 0.02, Friedman test). **j,** SVM classification performance for simple cloud-of-tones (evidence strength, −1 vs 1) and mixed cloud-of-tones (evidence strength, −0.6 or −0.1 vs 0.1 or 0.6). **k,** Quantification of SVM classification performance in (j), averaged over the cloud period (mixed, *P = 0.0178; simple, ***P = 1.91e-4, Student paired t test). **l,** Confocal images showing mPFC neurons expressing ArchT and optic fiber placement. **m,** Diagrams showing optic fiber placement in the mPFC of different mice. **n,** Psychometric curves for mice (n = 7 mice) in which the mPFC neurons were photo-inhibited (with laser) during the contextual period in two different sets of sessions in the absence of contextual cues (no-CX1 & no-CX2). **o,** Quantification of bias indices for the mice in **n** (n = 7, ns, P = 0.1527, Student paired t-test). **p,** Quantification of the change in bias index in the two different sets of sessions (n = 7, ns, P = 0.984; Student paired t-test). Psychometric curves are averaged over six sessions. Data are presented as mean ± s.e.m.

**Extended Data Fig. 3.**
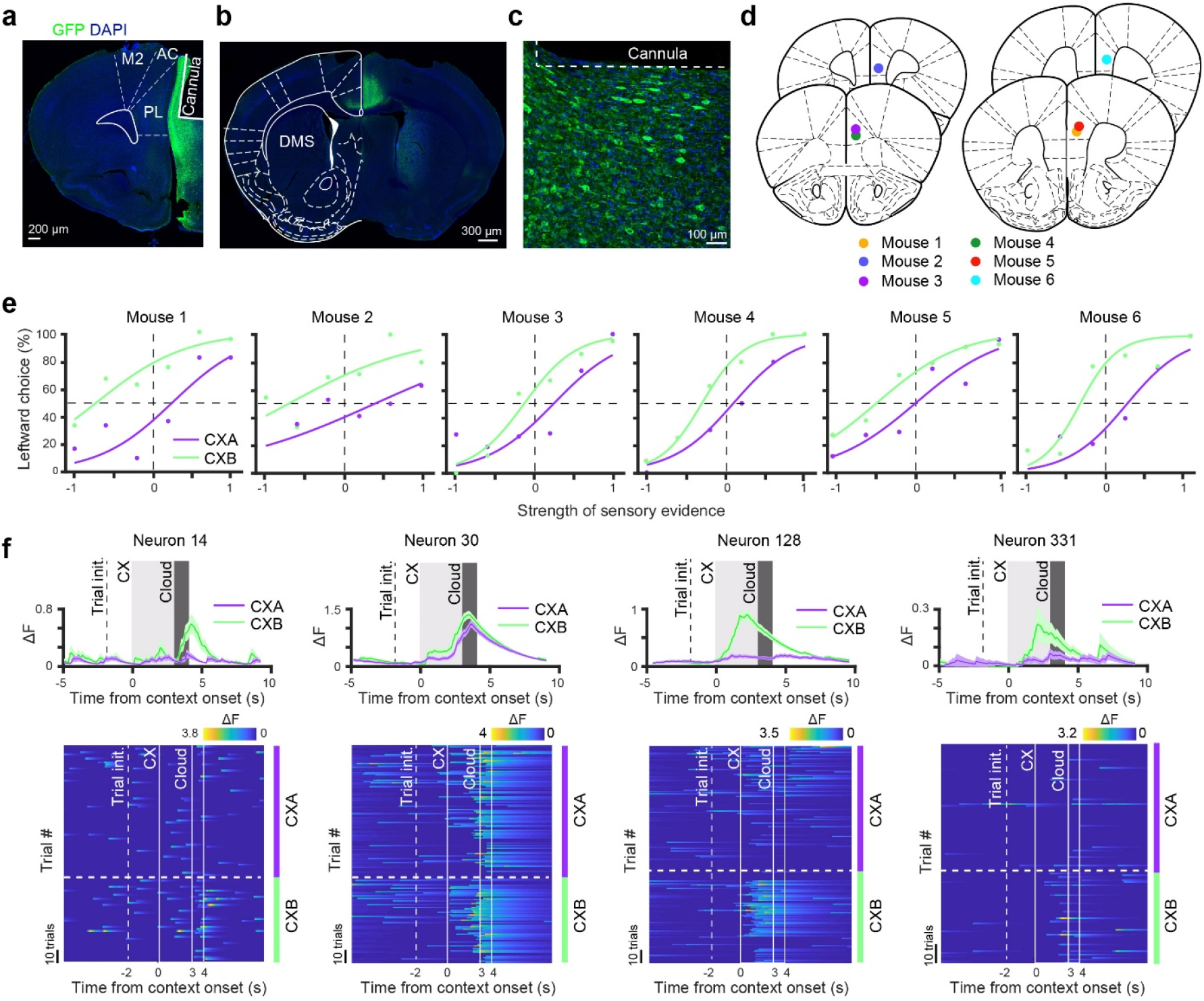
Imaging mPFC^DMS^ neuron activity during the CX2AC task. (**a-c**) Confocal images showing the expression of GCaMP6f and cannula placement in the mPFC (a), the GCaMP6f-labeled axon fibers in the DMS originating from the mPFC (b), and a higher magnification image of mPFC^DMS^ neurons expressing GCaMP6f (c). **d,** Diagrams showing cannula placement in the mPFC of different mice. **e,** Psychometric curves of individual mice during imaging. **f**, Responses of example mPFC^DMS^ neurons at the late learning stage. Top panel: average activity over all trials in each context. Bottom panel: heatmaps of trial-by-trial activity in each context. Data are presented as mean ± s.e.m.

**Extended Data Fig. 4.**
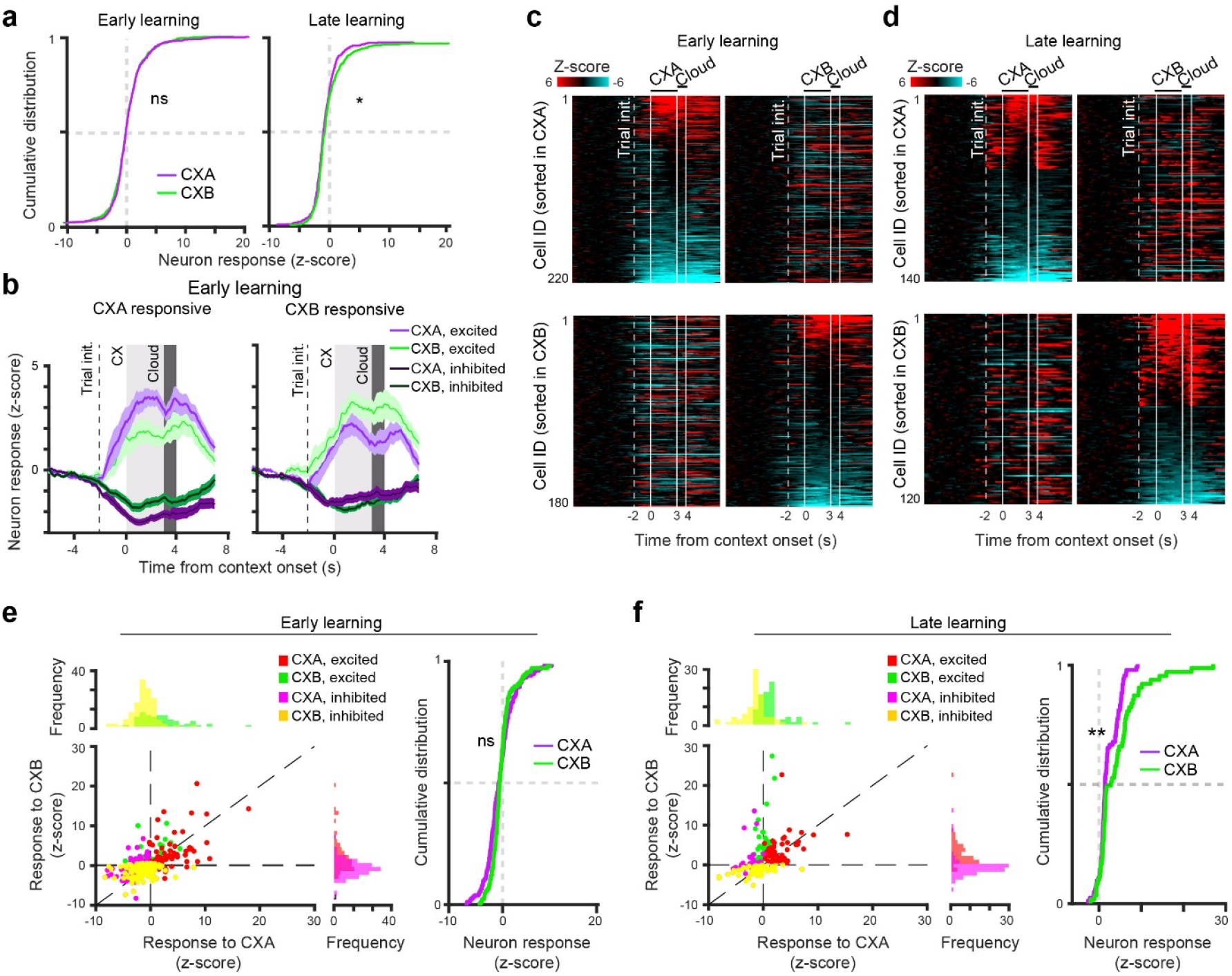
Characterization of mPFC^DMS^ neuron activity under different contexts in the CX2AC task. **a,** Cumulative distribution of the responses of all neurons to CXA and CXB presentations in the early (left) and late (right) learning stages (early, ns (nonsignificant), P = 0.55, late, *P = 0.013, Wilcoxon rank-sum test). **b,** Left: average activity of CXA-responsive (excited or inhibited) neurons in CXA trials and CXB trials. Right: average activity of CXB-responsive (excited or inhibited) neurons in CXA trials and CXB trials. Activity was acquired during the early learning stage. **c,** Heatmaps of the activity (z-scored) of individual neurons significantly (permutation test, P < 0.05) responding to CXA (top) and CXB (bottom) at the early stage of learning. Each row represents a neuron. Neurons are sorted according to their activity during CXA (top) or CXB (bottom). **d,** Same as (c), except that data were acquired during the late learning stage. **e,** Left panel: CXA-responses and CXB-responses of individual neurons shown in (c). Right panel: cumulative distribution of CXA-responses and CXB-responses of individual neurons shown in (c) (ns, P = 0.813, Wilcoxon rank-sum test). **f,** Left panel: CXA-responses and CXB-responses of individual neurons shown in (d). Right panel: cumulative distribution of CXA-responses and CXB-responses of individual neurons shown in (d) (**P = 0.0033, Wilcoxon rank-sum test).

**Extended Data Fig. 5.**
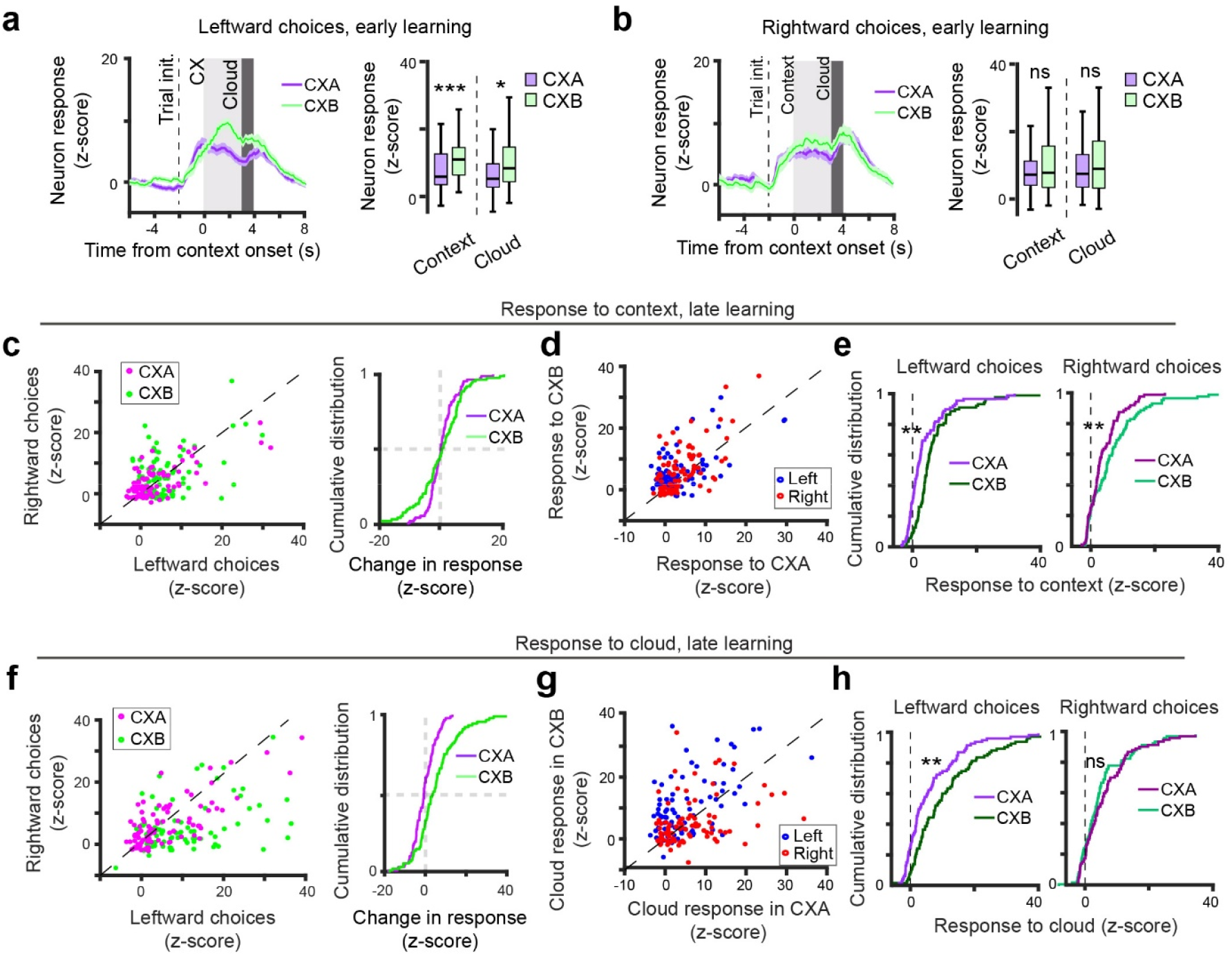
Characterization of mPFC^DMS^ neuron activity during leftward and rightward choices in the CX2AC task. **a,** Left panel: average activity of CXB-excited neurons in trials in which mice made leftward choices under CXA or CXB at the early stage of learning. Right panel: quantification of the activity during the context and cloud period (context, ***P = 0.001, cloud, *P = 0.022, Wilcoxon rank sum test). **b,** Same as (a), except that data were from trials in which mice made rightward choices (context, ns (nonsignificant), P = 0.357, cloud, ns, P = 0.562, Wilcoxon rank sum test). **(c-e)** Responses of individual neurons to contexts during the late stage of learning. **c,** Left panel: scatter plot of the responses of individual neurons during leftward and rightward choices in CXA or CXB trials (CXA, P = 0.24, CXB, P = 0.002, sign rank test). Right panel: cumulative distribution of response difference between leftward and rightward choices for individual neurons in each context. **d,** Scatter plot of CXA-response and CXB-response of each neuron during leftward or rightward choices. **e,** Cumulative distribution of CXA-responses or CXB-responses of individual neurons during leftward choices (left panel; **P = 0.0094, Wilcoxon rank sum test) and rightward choices (right panel; **P = 0.0082, Wilcoxon rank sum test). **(f-h)** Responses of individual neurons to cloud-of-tones stimuli during late stage of learning. **f,** Left panel: scatter plot of the responses during the leftward choices and rightward choices for individual neurons in CXA or CXB trials (CXA, P = 0.149, CXB, P = 2.01e-5, sign rank test). Right panel: cumulative distribution of response difference between leftward and rightward choices for individual neurons in each context. **g,** Scatter plot of the responses of each neuron in CXA trials and CXB trials during leftward or rightward choices. **h,** Cumulative distribution of the responses of individual neurons in different contexts during leftward choices (left panel; **P = 0.0015, Wilcoxon rank sum test) and rightward choices (right panel; ns, P = 0.3708, Wilcoxon rank sum test).

**Extended Data Fig. 6.**
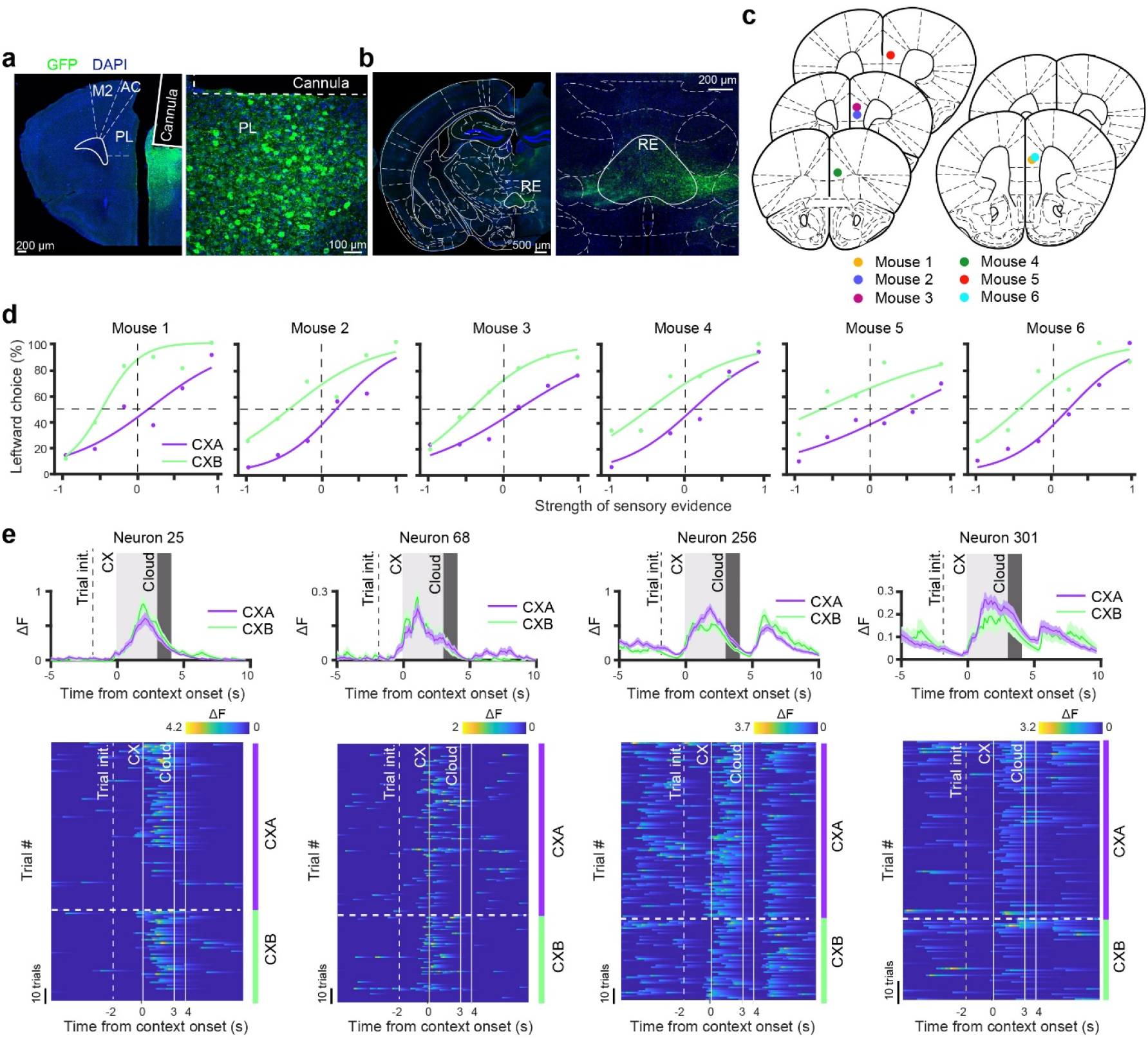
Imaging mPFC^VMT^ neuron activity during the CX2AC task. **a,** Confocal images showing the expression of GCaMP6f and cannula placement in the mPFC. On the right is a higher magnification image of the mPFC area in the image on the left, showing mPFC^DMS^ neurons expressing GCaMP6f. **b,** Confocal images showing the GCaMP6f-labeled axon fibers in the VMT area originating from the mPFC. On the right is a higher magnification image of the VMT area in the image on the left, showing the GCaMP6f-labeled axon fibers in the nucleus reuniens (RE) of the VMT. **c,** Diagrams showing the cannula placement in the mPFC of different mice. **d,** Psychometric curves of individual mice during imaging. **e**, Responses of example mPFC^VMT^ neurons at the late learning stage. Top panel, average activity over all trials in each context. Bottom panel, heat-maps of trial-by-trial activity in each context. Data are presented as mean ± s.e.m.

**Extended Data Fig. 7.**
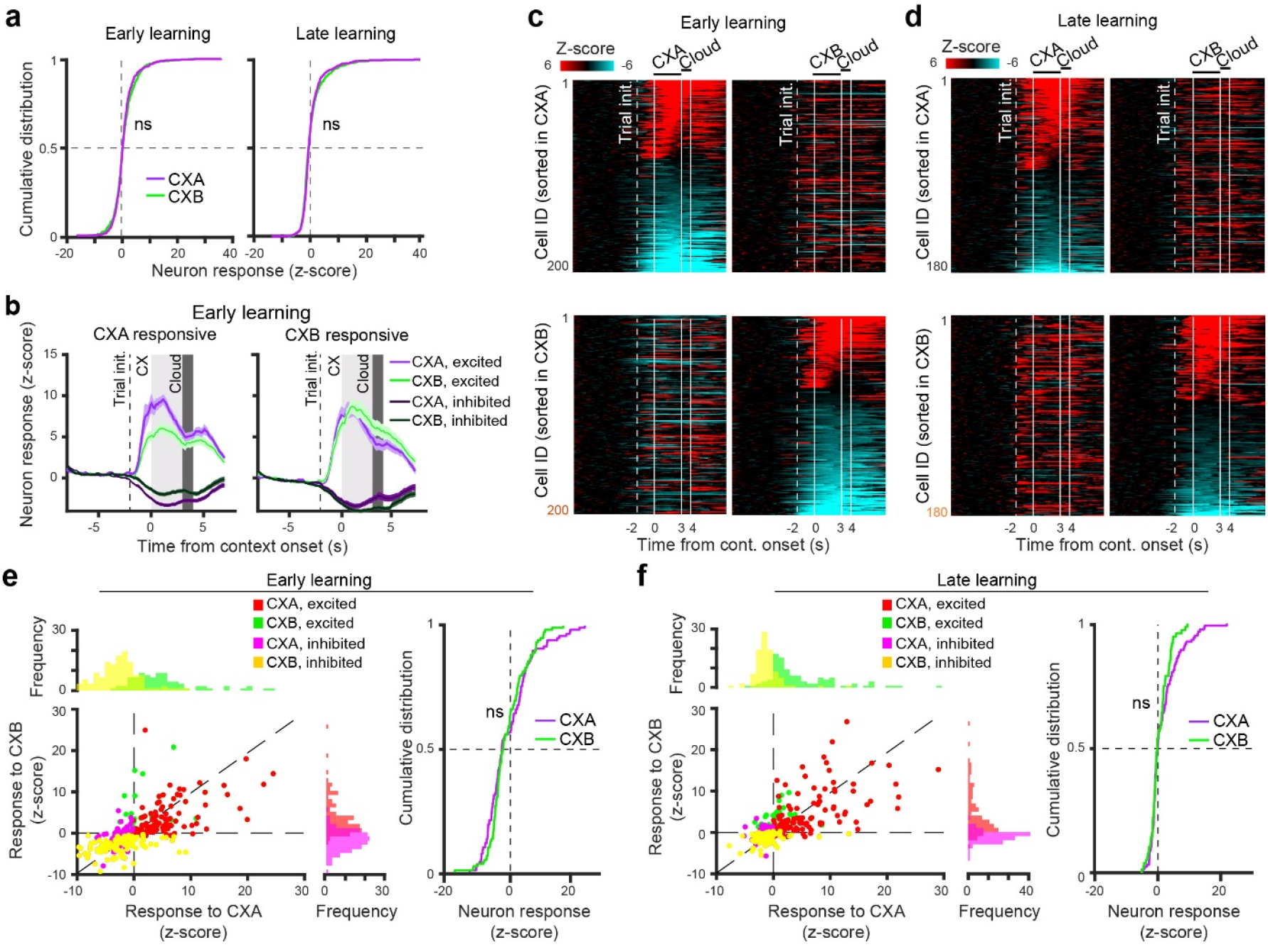
Characterization of mPFC^VMT^ neuron activity under different contexts in the CX2AC task. **a,** Cumulative distribution of the responses of all neurons to CXA and CXB presentations in the early (left) and late (right) learning stages (early, ns (nonsignificant), P = 0.41, late, ns, P = 0.97, Wilcoxon rank-sum test). **b,** Left: average activity of CXA-responsive (excited or inhibited) neurons in CXA trials and CXB trials. Right: average activity of CXB-responsive (excited or inhibited) neurons in CXA trials and CXB trials. Activity was acquired during the early learning stage. **c,** Heatmaps of the activity (z-scored) of individual neurons significantly (permutation test, P < 0.05) responding to CXA (top) and CXB (bottom) at the early stage of learning. Each row represents a neuron. Neurons are sorted according to their activity during CXA (top) or CXB (bottom). **d,** Same as (c), except that data were acquired during the late learning stage. **e,** Left panel: CXA-responses and CXB-responses of individual neurons shown in (c). Right panel: cumulative distribution of CXA-responses and CXB-responses of individual neurons shown in (c) (ns, P = 0.718, Wilcoxon rank-sum test). **f,** Left panel: CXA-responses and CXB-responses of individual neurons shown in (d). Right panel: cumulative distribution of CXA-responses and CXB-responses of individual neurons shown in (d) (ns, P = 0.424, Wilcoxon rank-sum test). Data are presented as mean ± s.e.m.

**Extended Data Fig. 8.**
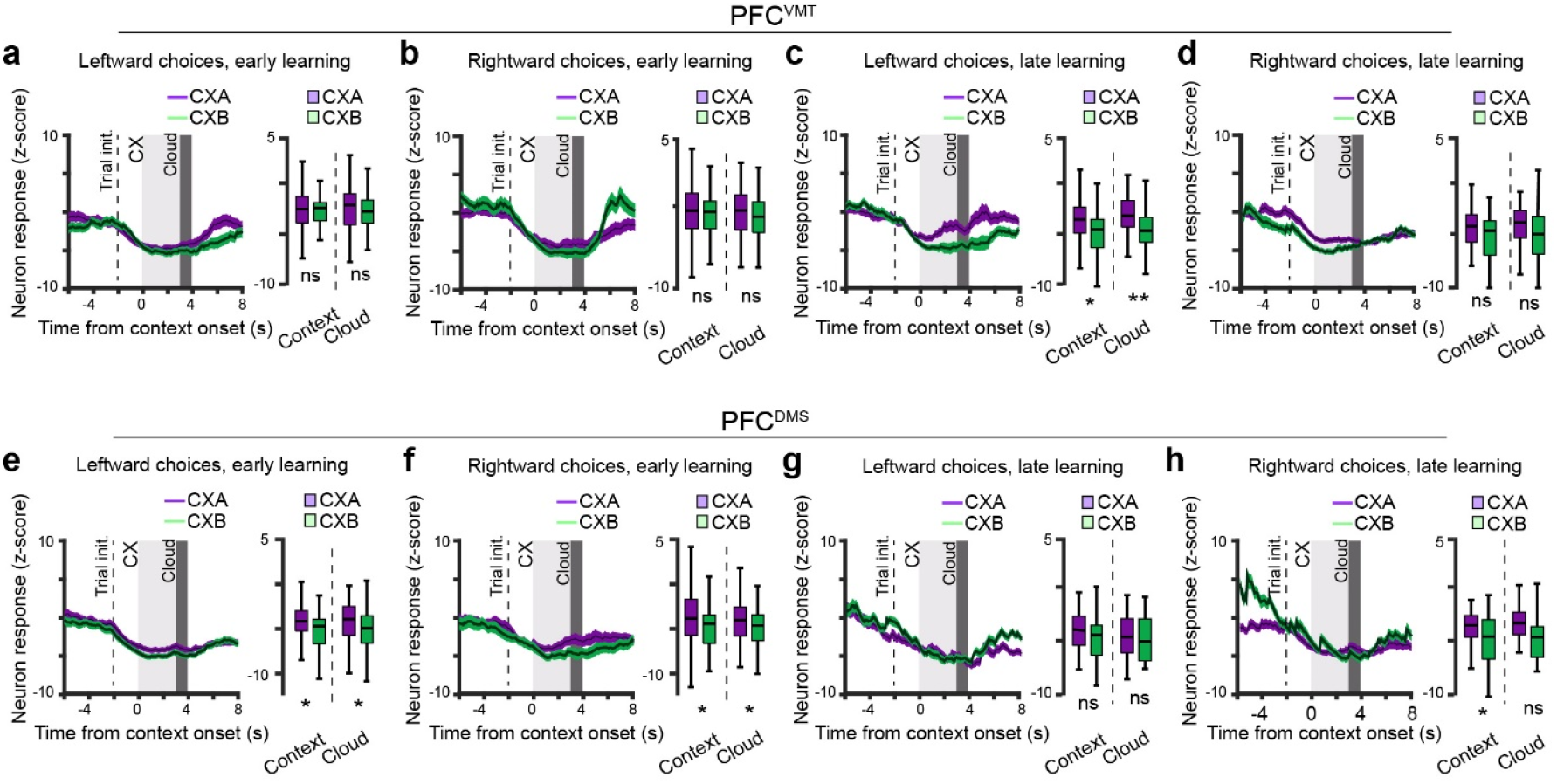
mPFC^VMT^ neurons, but not mPFC^DMS^ neurons, show decreased activity during context-dependent decisions. (**a-d**) Activity of mPFC^VMT^ neurons showing significant reductions (z-score < −3) in activity during the contextual period. **a,** Left panel: average activity in trials in which mice made leftward choices under CXA or CXB at the early stage of learning. Right panel: quantification of the activity during the context and cloud period (context, ns (nonsignificant), P = 0.469, cloud, ns, P = 0.142, Wilcoxon rank sum test). **b,** Same as (a), except that the analysis was for trials in which mice made rightward choices (context, ns, P = 0.246, cloud, ns, P = 0.178, Wilcoxon rank sum test). (**c, d**) Same as (a, b), respectively, except that data were acquired during the late stage of learning. **c,** Context, *P = 0.0143, cloud, **P = 0.002, Wilcoxon rank sum test. **d,** Context, ns, P = 0.069, cloud, ns, P = 0.189, Wilcoxon rank sum test. (**e-h**) Activity of mPFC^DMS^ neurons showing significant reductions (z-score < - 3) in activity during the contextual period. **e,** Left panel: average activity in trials in which mice made leftward choices under CXA or CXB at the early stage of learning. Right panel: quantification of the activity during the context and cloud period (context, *P = 0.031, cloud, *P = 0.02, Wilcoxon rank sum test). **f,** Same as (e), except that the analysis was for trials in which mice made rightward choices (context, *P = 0.015, cloud, *P = 0.026, Wilcoxon rank sum test). (**g, h**) Same as (e, f), respectively, except that data were acquired during the late stage of learning. **g,** Context, ns, P = 0.438, cloud, ns, P = 0.714, Wilcoxon rank sum test. **h,** Context, *P = 0.171, cloud, ns, P = 0.068, Wilcoxon rank sum test.

**Extended Data Fig. 9.**
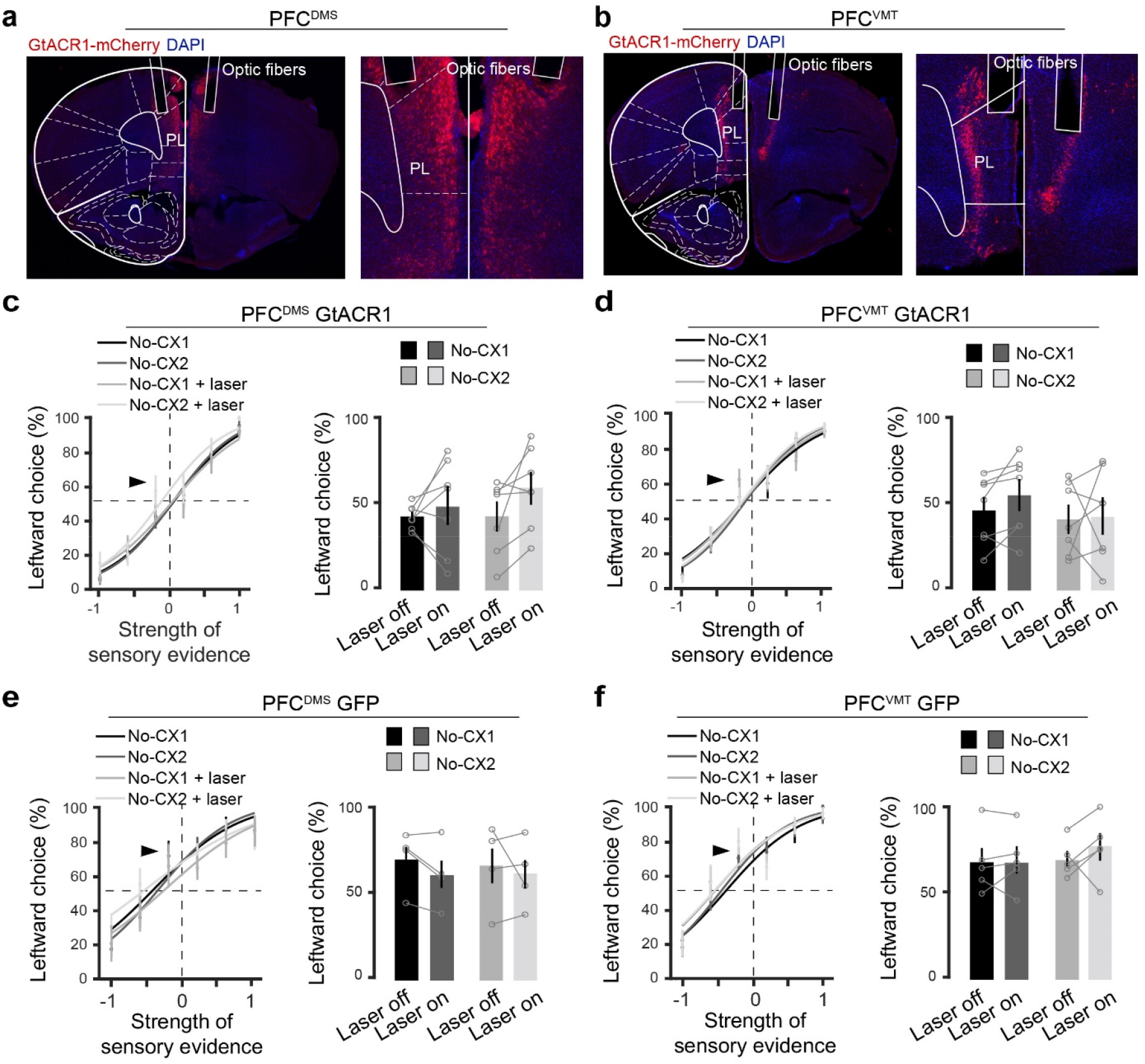
Histological verification and control experiments for the optogenetic inhibition of mPFC^DMS^ neurons and mPFC^VMT^ neurons. **a,** Confocal images showing the expression of GtACR1 in mPFC^DMS^ neurons and optic fiber placement in the mPFC. On the right is a higher magnification image of the mPFC area in the image on the left. Note that mPFC^DMS^ neurons are located in layer 2/3. **b,** Confocal images showing the expression of GtACR1 in mPFC^VMT^ neurons and optic fiber placement in the mPFC. On the right is a higher magnification image of the mPFC area in the image on the left. Note that mPFC^VMT^ neurons are located in layer 5/6. **c,** Left panel: psychometric curves of mice in the CX2AC task, in which mPFC^DMS^ neurons expressed GtACR1. In randomly interleaved trials, the mPFC^DMS^ neurons were photo-stimulated during the period between trial initiation (licking at the center spout) and the onset of the cloud-of-tones, but in the absence of any contextual cues. This procedure was repeated in two different sets of sessions without contextual cues (no- CX1 & no-CX2). Right panel: quantification of leftward choices in the trials with cloud-of- tones of “−0.1” evidence strength (indicated by the arrowhead on the psychometric curves) (F(1,7) = 0.7, P = 0.44, two-way ANOVA with repeated measures). **d,** Same as (c), except that data were from mice in which mPFC^VMT^ neurons expressed GtACR1 (F(1,7) = 0.565, P = 0.44, two-way ANOVA with repeated measures). **e,** Same as (c), except that data were from mice in which mPFC^DMS^ neurons expressed GFP (F(1,3) = 0.872, P = 0.91, two-way ANOVA with repeated measures). **f,** Same as (c), except that data were from mice in which mPFC^VMT^ neurons expressed GFP (F(1,3) = 3.88, P = 0.08, two-way ANOVA with repeated measures).

